# NeuroDISK: An AI Approach to Automate Continuous Inquiry-Driven Discoveries in Neuroimaging Genetics

**DOI:** 10.1101/2025.02.14.638360

**Authors:** Daniel Garijo, Qifan Yang, Hernán Vargas, Shruti P. Gadewar, Kevin Low, Varun Ratnakar, Maximiliano Osorio, Alyssa H. Zhu, Agnes McMahon, Yolanda Gil, Neda Jahanshad

**Affiliations:** Information Sciences Institute, University of Southern California, Marina del Rey, California, USA; Ontology Engineering Group, Universidad Politécnica de Madrid, Madrid, Spain; Laboratory of Brain eScience, Mark and Mary Stevens Neuroimaging and Informatics Institute, Keck School of Medicine of USC, University of Southern California, Marina del Rey, California, United States; Department of Neurology, Keck School of Medicine of USC, University of Southern California, Marina del Rey, California, USA; Department of Biomedical Engineering, Viterbi School of Engineering, University of Southern California, Marina del Rey, California, USA

## Abstract

Collaborative and multi-site neuroimaging studies have greatly accelerated the rate at which new and existing data can be aggregated to answer a neuroscientific question. New research initiatives are continuously collecting more data, allowing opportunities to refine previous published findings through continuous and dynamic updates. Yet, we lack a practical framework for researchers to systematically, automatically, and continuously update published findings. We developed NeuroDISK, an automated artificial intelligence based framework that: 1) performs automated and inquiry-driven analyses, and 2) continuously updates these analyses as new data becomes available. NeuroDISK was evaluated using published results from the ENIGMA consortium’s work on the genetic architecture of the cerebral cortex. We incorporate both meta-analysis and meta-regression options to showcase our framework on the effect of specific genotypes and moderators on select brain regions. Initial NeuroDISK meta-analysis results replicate the original publication, and we show result updates after adding new data. The NeuroDISK framework can be generalized for users to define question(s), run corresponding workflow(s) and access results interactively and continuously.

## 1. Introduction

Sample sizes for studies involving human subjects are often limited by the costs of collecting data. Unfortunately, in these studies, variables of interest that have small to moderate effect sizes have been particularly susceptible to spurious findings that do not replicate. Recent efforts have recognized these concerns in neuroimaging-heavy fields of psychological and neurological sciences(Button et al., 2013),(Boekel et al., 2015),(Bowring et al., 2019),(Poldrack et al., 2020),(Hodge et al., 2020), and even more specifically, neuroimaging genetics(Medland et al., 2014; Smith & Nichols, 2018). A driving factor in this reproducibility and replication crisis has been a lack of sufficiently well-powered studies,(Button et al., 2013; Ioannidis, 2005) so larger scale efforts are of growing interest.

Merging data from multiple sources has paved a way for larger sample sizes and reproducible findings. Individual multi-site studies such as the disorder-specific Alzheimer’s Disease Neuroimaging Initiative (ADNI)(Weiner et al., 2010) and the population-based UK Biobank(Miller et al., 2016) have aimed to collect both neuroimaging and genetic data across individuals from multiple locations for larger and more efficient data collection efforts. Multi-study consortia such as the Enhancing NeuroImaging Genetics through Meta Analysis (ENIGMA) consortium(Thompson et al., 2020) have also been established to coordinate analyses and pool data across tens of thousands of brain imaging datasets from hundreds of independent studies around the world. ENIGMA has over 35 active working groups with targeted clinical, biological or methodological interests. These working groups, including one dedicated to neuroimaging genetics, pool data from independent existing studies from around the world. Consortia that are built on existing and available data resources not only ensure large sample sizes for well-powered analyses, but also include diverse samples and heterogeneous study designs to allow for robust and generalizable findings. Multi-study efforts in ENIGMA ensure adequate sample sizes for genome-wide association studies (GWAS) on brain imaging derived traits(Medland et al., 2022) and have led to the identification of numerous genetic variants that influence brain structure through some of the largest studies to date.

As ENIGMA studies are being conducted, new neuroimaging genetics initiatives and new studies with relevant data continue to be funded and collected. These efforts may eventually be incorporated into other mutli-site and multi-study initiatives, further empowering larger and more representative studies. For example, in 2012 a multi-study GWAS of hippocampal volume was published by the ENIGMA consortium(Stein et al., 2012). Interest soon grew in the consortium, and a follow up study that again included a GWAS of the hippocampal volume was conducted with nearly twice the sample size(Hibar et al., 2015). The same inquiry was then posed jointly by ENIGMA and another multi-study consortium, CHARGE (Cohorts for Heart and Aging Research in Genomic Epidemiology), more than doubling the sample size once again(Satizabal et al., 2019). These studies have iteratively shown the progression of insights into the genetic architecture of regional brain volumes, including that of the increasing confidence in the effect of a genetic locus on chromosome 12 on the volume of the hippocampus (**Figure 1**). This was the only locus to show a genome-wide significant effect in the first analysis, and confidence of this finding only grew in subsequent analyses. The evolution of scientific knowledge is captured by repeatedly making the same inquiry, yet with more or different data. *Artificial intelligence (AI) systems that can automatically generate these updates as more data become available, with minimal human intervention, can greatly facilitate research efficiency and accelerate scientific advances*.

**Figure 1:**
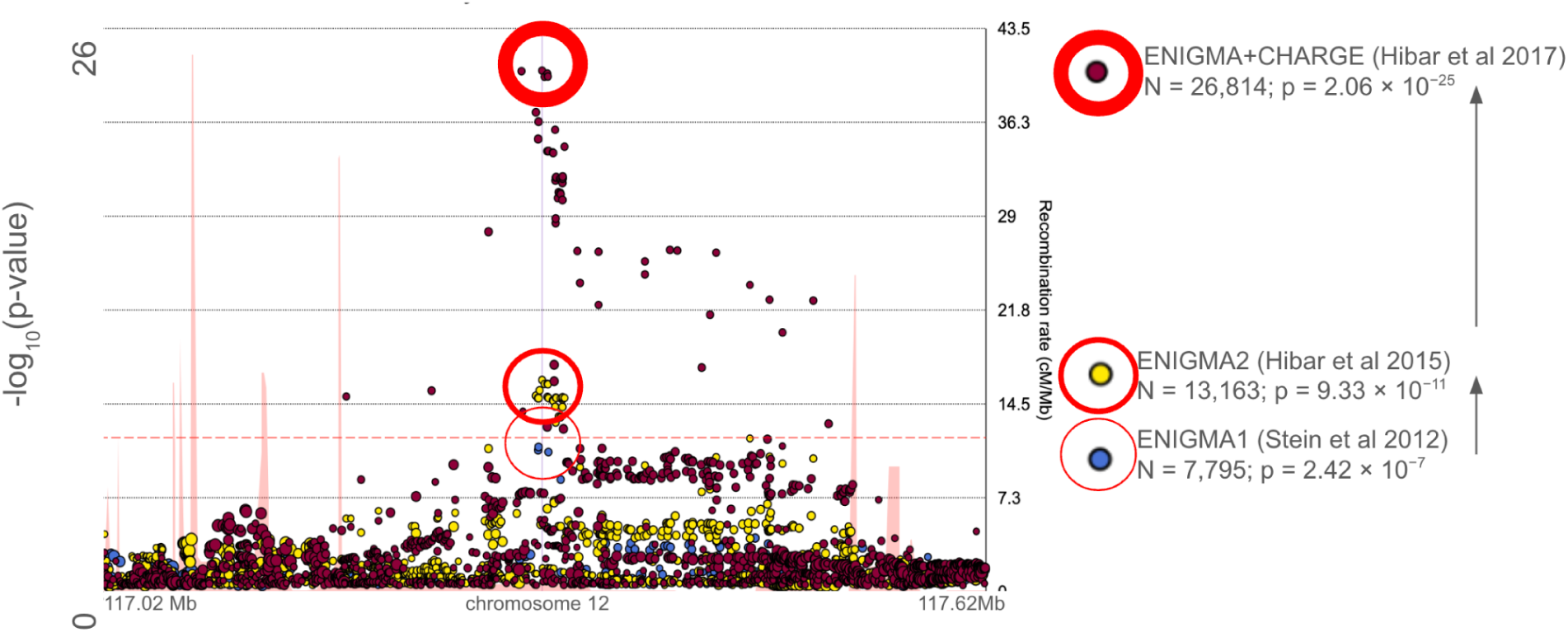
As more and more data become available for collaborative efforts, including neuroimaging genetics studies, a hypothesis can be tested with more data to increase confidence. In the above graphic, we show the evolution of results related to the genetic associations with MRI-derived hippocampal volume. The most significant genetic association with hippocampal volume in the first ENIGMA consortium study (ENIGMA1)(Stein et al., 2012), which performed a genome-wide association study (GWAS) meta-analysis of the hippocampus and the brain’s intracranial volume with data for almost 8,000 individuals across 17 cohorts, was in a locus on chromosome 12. In the second ENIGMA GWAS (ENIGMA2)(Hibar et al., 2015), which evaluated hippocampal volume along with 6 other structures in a pooled sample of over 13,000 individuals from 28 cohorts, the same locus (middle panel) emerged as significant with greater confidence (smaller p-value). When the results of ENIGMA2 were then meta-analyzed with those from the CHARGE consortium in an extended analysis of 9 subcortical structures using a total sample of over 26,000 individuals from 46 discovery cohorts (ENIGMA+CHARGE) (Satizabal et al., 2019), again the same genetic locus showed significant association with the hippocampal volume with greater confidence. The thickness of the red circle indicates the strength of the association, highlighting the greater significance as the dataset is expanded (bottom to top). Specifically, blue points reflect the ENIGMA1 study from 2012(Stein et al., 2012); yellow points correspond to the ENIGMA2 study from 2015(Hibar et al., 2015), and burgundy points represent the ENIGMA+CHARGE joint analysis from 2017(Hibar et al., 2017) (thickest red circle). We show the extent to which the significance changed from study to study. The same locus was then separately shown to be significant in independent data from over 8,000 individuals in the UK Biobank dataset(Elliott et al., 2018), which had not been used in any of the initial three ENIGMA or ENIGMA-CHARGE publications.

These AI systems would conduct continuous monitoring to detect new data and re-execute analyses to update findings. Intelligent automation can further be used to interrogate the data as more and more of the population becomes represented in the available data. For example, if the support for an association becomes stronger or weaker once more data is added and the sample becomes more diverse, an intelligent system may be able to identify aspects of the populations that were driving the changes in the association strength. We have previously found that the mean age of a study’s participants may drive key associations between neuroimaging and genetic markers, including the association between hippocampal volume and one of the strongest genetic risk factors for Alzheimer’s disease and related dementias(Brouwer et al., 2022; Garijo et al., 2019); here associations were only identified in cohorts, or datasets, of studies where the average age was over 60 years.

This work presents a two-fold AI approach to: 1) perform automated and inquiry-driven analyses, and 2) continuously update these analyses as new data becomes available. We have designed and implemented NeuroDISK, an AI system built on the DISK framework for assessing hypotheses evolution.(Garijo et al., 2017; Gil et al., 2016) NeuroDISK currently focuses on queries using a pilot set of neuroimaging genetics tasks given a structured data ontology and specific analytical workflows with added constraint reasoning.

Our work demonstrates automation of inquiry-driven data analysis in science, demonstrating that AI can reason about data and methods to automate this process for neuroimaging. AI has been investigated for a long time to automate data analysis and discovery processes in other fields of science (Bradshaw, Gary L., Zytkow, Jan M., 1988). Recent work uses workflows and reasoning to create automated machine learning systems (De Bie et al., 2022) and automated statisticians (Steinruecken et al., 2019). Other work has investigated the use of AI-based algorithms to accelerate experiment design and other scientific tasks. Rodriguez and colleagues (Rodríguez et al., 2024) demonstrate the use of AI to accelerate optical microscopy experiments over traditional numerical optimization techniques. Krenn and colleagues (Krenn et al., 2021) describe Theseus, a system that automates the design of quantum optics experiments based on a graph of known experiments. Furthermore, Erps and colleagues (Erps et al., 2021) describe an approach to automate the experimental design of formulations for mixed polymer formulations for additive manufacturing through multiobjective optimization.

AI automation of science processes has led to many advances in chemistry and materials science. In addition to the design and planning of experiments, there are approaches that add an execution component through AI and robotics (Angelopoulos et al., 2024). For example, Szymanski and team (Szymanski et al., 2023) introduce A-lab, a system to synthesize novel materials, specifically solid-state synthesis of inorganic powders. A-lab uses LLMs to propose new experiments, machine learning to optimize them, and a robotic platform to execute them. Pyzer-Knapp and colleagues (Pyzer-Knapp et al., 2022) describe an AI prototype for materials discovery, including natural language tools to extract information from the literature, simulators to optimize molecule choices, and a robotic chemistry platform to execute the experiments.

The use of large language models can accelerate research tasks, from literature search to code generation. For example, Coscientist (Boiko et al., 2023) was developed as a proof-of-concept system that designs and plans Suzuki and Sonogashira chemical reaction experiments and generates code that can be executed in cloud laboratories. In other work, Lu and colleagues (Lu et al., 2024) describe a system that uses large language models to generate ideas for papers, design the experiment, find the data and code needed to execute the experiment, and then write the article. The articles generated are useful for exploring new ideas and their feasibility, however the articles are not guaranteed to contain correct statements.

Related work in neuroimaging has focused on reproducibility of data analysis. This includes using software containers to ensure accurate replication (Renton et al., 2024), articulating requirements for data repositories to support reproducibility (Wagner et al., 2022), and infrastructure to access and integrate data and tools (Poline et al., 2023). Our work builds on these kinds of efforts by assuming the data and tools are shared, and extends them to demonstrate the use of AI to automate neuroimaging analyses.

We demonstrate the value of NeuroDISK by using and building on published data from the large-scale multi-study GWAS meta-analysis of MRI-derived cortical structure (Grasby et al., 2020). We catalog the published data and meta-data from all cohort studies that contributed to that work, and automate the statistical meta-analysis to demonstrate a successful replication of the available original findings. NeuroDISK then identifies newly cataloged data that match the requirements for the specific neuroimaging genetic inquiry and incorporates it into the analysis, ultimately updating the findings of the original paper. We further demonstrate the capabilities of NeuroDISK by asking additional questions about the data beyond what was originally proposed; in this demonstration, we inquire whether study specific effects are driven by particular aspects of the study cohort, in particular, mean age.

The contributions of this paper include:

1. A new AI approach to scientific problem solving designed to capture the strategies that a scientist follows to answer a question or test a hypothesis, including finding data, analyzing it, and extracting findings.
2. A novel concept of lines of inquiry that can automate hypothesis-driven discovery processes using AI knowledge representation and reasoning techniques that include ontologies, constraint reasoning, and workflows.
3. An implementation of this approach in NeuroDISK, an extension of the general domain-independent DISK framework (Gil et al., 2017), is extended with hypothesis ontologies and lines of inquiry for multi-site neuroimaging genetics.
4. A demonstration of this framework as a reproduction of a published paper, with explicit questions and hypotheses driving the system, and an extension of the published results to demonstrate its use for continuous updates.

In the following sections, we motivate the requirements for achieving the desired capabilities of automation and continuous updates, and describe the approach and its implementation.

## 2. Methods and Materials

NeuroDISK is designed to mimic how human scientists pursue a scientific question or hypothesis. Generally scientists pose questions or hypotheses and consider different approaches to answer their questions, which typically involve finding relevant data, and identifying appropriate analytical methods. For example, if MRI data is available it would be analyzed using computational imaging methods, while genomics data would be analyzed using genomics workflows. Finally, the results would be consolidated and placed in context.

NeuroDISK demonstrates how to automate this inquiry-driven discovery process for neuroimaging genomics. NeuroDISK uses AI representations and reasoning to test and revise hypotheses based on automatic analysis of scientific data repositories that grow over time (**Figure 2**). Two key features of NeuroDISK are: 1) *Inquiry-driven automated analysis*: Given an input hypothesis or scientific question, NeuroDISK is able to automatically search for relevant data in shared repositories and apply appropriate methods to test it; 2) *Continuous automated updates of findings*: NeuroDISK checks if new data becomes available so it can reconsider prior analyses and revise its findings.

**Figure 2.**
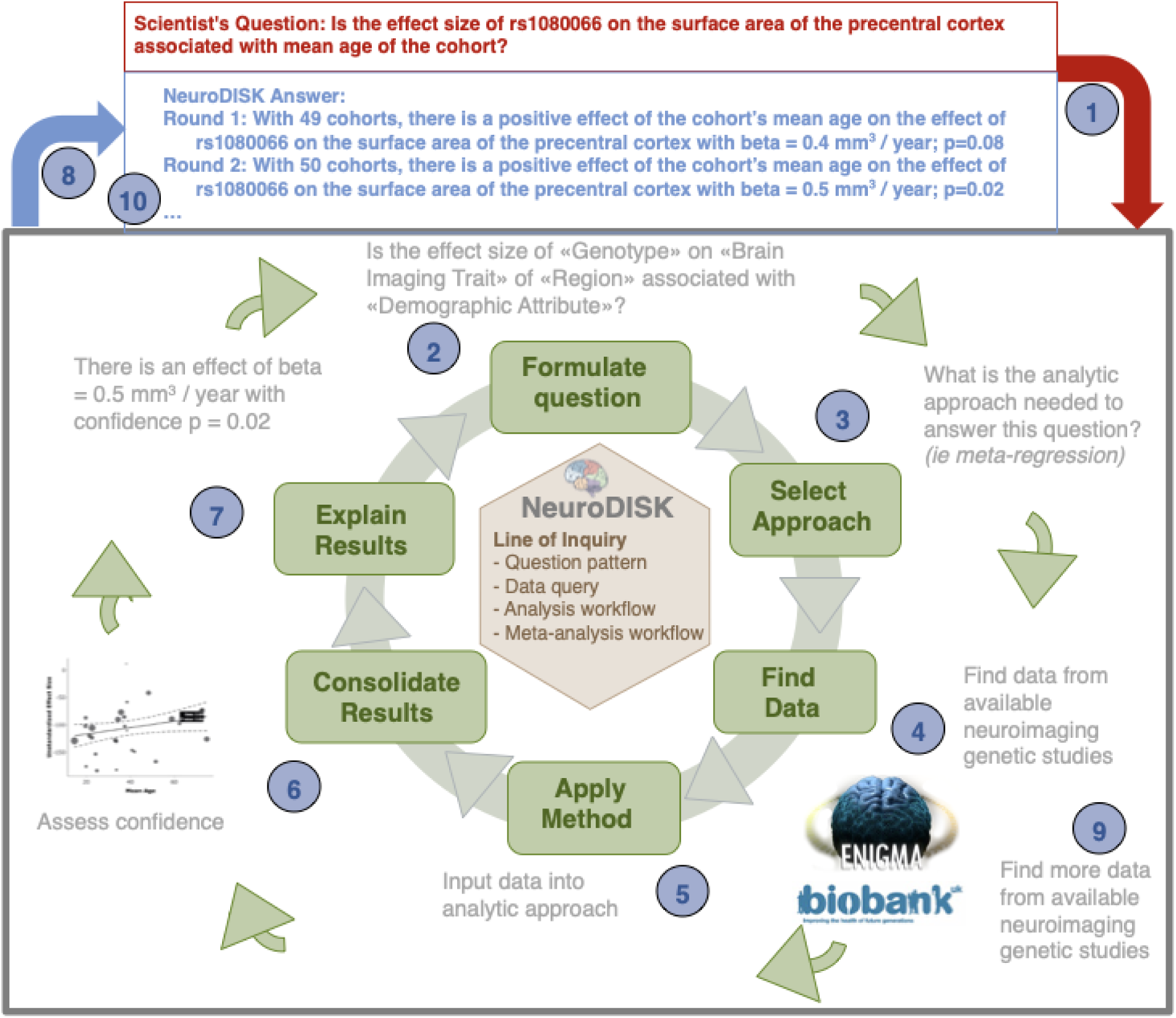
NeuroDISK is designed to automate the processes that scientists follow to answer questions using existing datasets. A scientific question (1) is mapped into a structured form that is machine-readable (2) that enables NeuroDISK to select a general approach (3) to answer that question. The approach typically involves formulating a query that will access existing data sources to find relevant datasets (4), setting up and running analyses for the data available (5), consolidating the results for different datasets (6), and explaining (7) and presenting (8) the findings. When new data become available (9), NeuroDISK revisits the original question and re-runs its analyses so that the findings can be updated (10).

### 2.1 Scientific methods as lines of inquiry

NeuroDISK uses a *line of inquiry* (LOI) to represent the approach that a scientist would follow to pursue a general type of question in their discipline, including steps to get data from a shared data source and to analyze it with computational workflows (**Figure 3**). An LOI has several key components:

- *Hypothesis or question template*: a general question containing *variables* that are matched against the specific question posed by a scientist.
- *Data query template*: indicates how a data source should be queried in order to obtain data that is relevant to the question. The data query template includes the variables that appear in the question template, as well as additional variables that characterize the data requirements in detail for the approach being pursued.
- *Workflows*: specify the multi-step methods to be used to analyze the data retrieved. The workflows use variables to indicate how to take the data retrieved as input. Workflows also represent the computational steps of the analysis method.
- *Meta-workflows*: specify the meta-method to combine results from multiple analyses (i.e., multiple workflow executions) in order to derive an answer to the user’s question. Meta-workflows combine results from other workflows and may generate overall statistics such as an effect size, confidence interval, p-value, or a refinement of the original question or hypothesis, as well as visualizations of results and findings.

**Figure 3.**
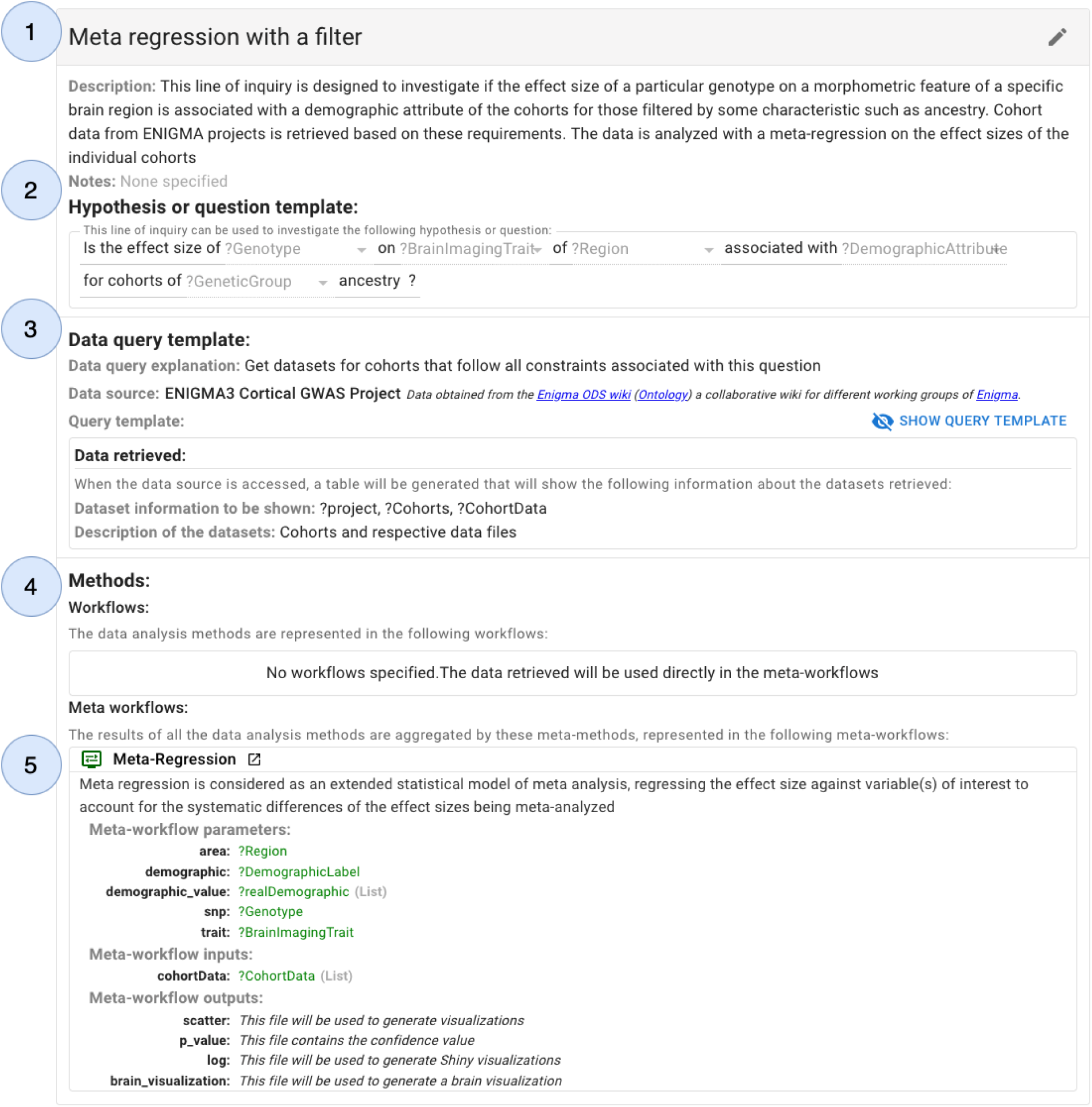
NeuroDISK uses Lines of Inquiry (LOIs) to represent the approach that a scientist would follow to answer different types of questions. An LOI consists of: (1) documentation about the LOI, which can include literature citations that introduce the approach; (2) a question template that will be matched against the user’s question; (3) a data query template that specifies how to retrieve data that is relevant to answering the question; (4) workflows that specify how to analyze the data retrieved; and (5) meta-workflows that indicate how to combine the results of all the workflows executions. In this example, an LOI is set up to investigate if the effect size of a particular genotype on a specific brain region is associated with a demographic attribute of the cohorts, while allowing the cohorts to be filtered by another attribute (in this case genetic ancestry). This process is a meta-regression on the effect sizes of the individual cohorts retrieved from the data sources.

The individual LOI components are described in more detail in this section.

Figure 3 illustrates the main components of an LOI for investigating if the effect size of a particular genotype on a specific brain region is associated with a demographic attribute of the cohorts, while allowing the cohorts to be filtered by another attribute (in this case genetic ancestry). This process is a meta-regression on the effect sizes of the individual cohorts retrieved from the data sources.

NeuroDISK captures knowledge in machine-readable representations that use Semantic Web standards, in particular the W3C Resource Description Framework (RDF)(*RDF 1.1 Concepts and Abstract Syntax*), the W3C Web Ontology Language (OWL)(Dean & McGuinness), and the W3C Semantic Protocol and RDF Query Language (SPARQL)(Harris et al., 2013).

### 2.2 Specifying inquiries: hypotheses and questions

NeuroDISK is an inquiry-driven system that expects users to provide a structured hypothesis or scientific question that will drive its reasoning and data analysis. Users are guided through pre-defined *question templates* and select one to specify an inquiry in a structured, machine-readable representation. Once that selection is made, the user’s question is matched against the query template of the available LOIs, which will trigger an appropriate one.

As our running example, we use results published as part of the 2020 ENIGMA cortical structure GWAS paper(Grasby et al., 2020). The paper, among its top findings, found an association between the genetic variant rs108066 on the surface area of the precentral gyrus after meta analyzing genome-wide association results from 48 discovery cohorts of individuals of European ancestry, replicated the findings in the UK Biobank, and generalized the findings in datasets of non-European individuals. We use this particular finding (i.e., the genetic variant association with regional brain area) as our running example. Using this example, we continue to build on the result by including independent data and formulate a novel scientific question to be asked of the available data. The novel question presented in NeuroDISK is to use available meta-data and individual cohort results to determine whether the strength of the particular genetic variant’s (rs108066) association with the specific regional brain metric (surface area of the precentral cortex) is associated with a demographic property of the cohorts (the mean age); we further allow filtering of the cohorts by specific properties (genetic ancestry). The user would start by selecting from a set of question templates. In this example, they would select: “*Is the effect size of «Genotype» in «Brain Imaging Trait» of «Region» associated with «Demographic Attribute» for cohorts of «Genetic Ancestry»?*”, with the brackets indicating variables. Then the user would be offered choices to select the desired variable values. The result would be the *user question*.

Question templates are expressed as a machine-readable *question pattern* that consists of RDF triples of the form {subject predicate object}. For our running example, this would be an RDF triple in a question pattern:

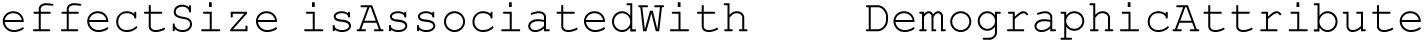

To specify question patterns, we follow a principled approach by using a Scientific Questions Ontology (SQO) that organizes question templates and variables(Garijo et al., 2023). This allows us to relate different scientific questions and to connect the user questions to LOIs. The SQO ontology is extended to create a Scientific Domain Ontology (SDO) with additional terms from any new domain that can be used to specify question variables and choices. The data source also uses a metadata ontology to describe datasets, which we refer to as the Data Source Ontology (DSO). DSO should be designed to extend SDO, or at least should be well mapped to it in order to support the specification of the data queries needed to support the anticipated user queries. In NeuroDISK, the SDO is the SDO-ENIGMA ontology and the DSO is the Organic Data Science (ODS)-ENIGMA ontology, which will be described in detail in **Section 2.3**.

Figure 4 illustrates the key concepts in the SQO and how they are used to create questions in the SDO in NeuroDISK. We describe these SQO concepts first and then provide examples. Key SQO concepts include:

- <u>Question Category</u> which helps organize all the question templates into broad types of scientific inquiries such as association, counterfactual, prediction, etc.
- <u>Question</u>: The class of user questions. Each question class includes:

- A Template, which consists of a text in natural language containing slots for question variables to be specified by a user.
- <u>Variables</u> that are filled by the user to express their question. In the example above, a variable can be the demographic which can be used to express user questions about age.
- <u>Constraints</u>, which are logical expressions that represent the valid combinations of variable values. These are used to ensure that candidate questions make scientific sense.
- A Pattern, which is a collection of RDF triples that combines the pattern fragments for all the question variables as described below.
- <u>Question Variable</u>: Each question variable is represented by:

- Variable Name:Denotes the name of the variable, e.g «Demographic»
- <u>Option</u>: Indicates the values that the question variables can take. The possible values can be indicated in several ways:

▪ <u>Static options</u>: A list of options pre-defined on the SDO. In our running example, the SDO-ENIGMA ontology has a variable «BrainImagingTrait» that has options ‘Surface Area’, ‘Thickness’, etc.
▪ <u>Dynamic options</u>: A list of options generated at run time by querying the data source, which would return terms from DSO. This query will use the question’s Constraints. For example, to generate options for a «Region» variable, the DSO for the data source would return the brain regions that are covered in the datasets available, such as the cortical regions used in this work: ‘Precentral’, ‘Insula’, etc.
▪ <u>User input options</u>: Allows the user to specify new variable values directly through the user interface as free form input. Users can only do this if they are very familiar with the domain and the datasets.
- <u>Min/max Cardinality</u>: Minimum/maximum number of options that can be selected by the user for the variable. By default each variable accepts and requires only one option (cardinality of 1).
- Pattern Fragment: The semantic expression (RDF triple) about this particular question variable, and that is part of the question pattern for a question.

**Figure 4.**
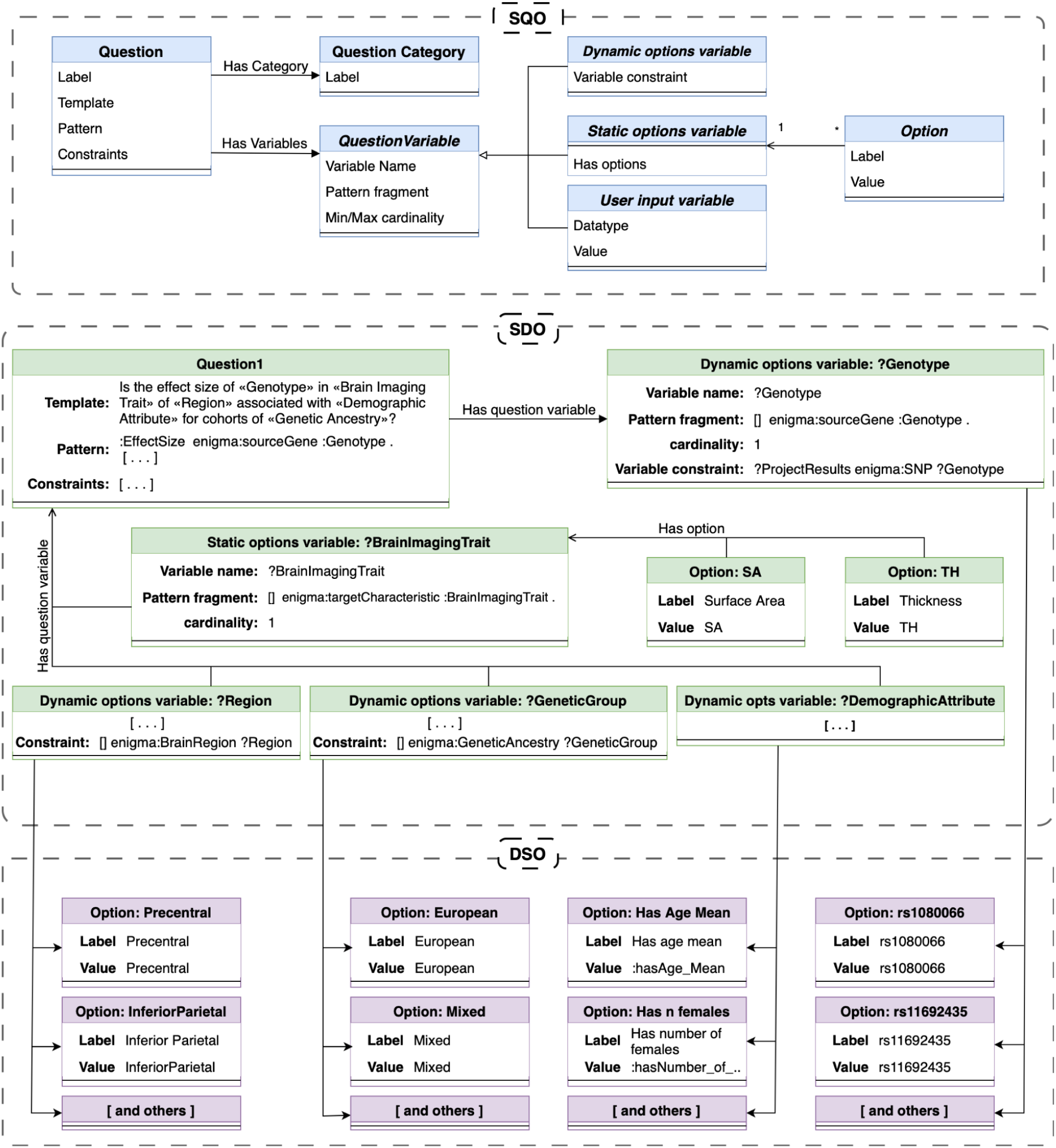
An overview of the ontologies in NeuroDISK. The Scientific Questions Ontology (SQO) at the top illustrates the main concepts and terms. Question templates in NeuroDISK have variables that users will fill out based on the options specified in the Scientific Domain Ontology (SDO) with additional options that are dynamically retrieved from the data source and expressed in a Data Source Ontology (DSO), as shown in the lower section of the figure. In NeuroDISK, the SDO is the SDO-ENIGMA ontology and the DSO is the ODS-ENIGMA ontology. Each ontology is grouped within a dashed box, where concepts are shown in different colors with labeled arrows indicating concept properties, and arrows across ontologies indicating subclasses.

Question patterns are expressed as a set of triples using terms from the SQO as well as the SDO. For our running example, the question template has the following question pattern:

|  |  |  |
| --- | --- | --- |
| sdo:effectSize | sdo:sourceGene | sdo:Genotype |
| sdo:effectSize | sdo:targetCharacteristic | sdo:BrainImagingTrait |
| sdo:effectSize | sdo:targetCharacteristic | sdo:Region |
| sdo:effectSize | sdo:isAssociatedWith | sdo:DemographicAttribute |
| sdo:effectSize | sdo:collectedFrom | sdo:GeneticAncestry |

When a user specifies their question and chooses variable values, the user question pattern is instantiated with the corresponding variable values (using the ‘ods:’ prefix for the values extracted dynamically from the data source and that are in the ODS-ENIGMA ontology). For example, when a user chooses *rs1080066* for Genotype, *Precentral Cortex* for Region, and *Surface Area* for BrainImagingTrait, these are the triples that represent those choices in the user question:

|  |  |  |
| --- | --- | --- |
| sdo:effectSize | sdo:sourceGene | ods:rs1080066 |
| sdo:effectSize | sdo:targetCharacteristic | ods:SurfaceArea |
| sdo:effectSize | sdo:targetCharacteristic | ods:PrecentralCortex |
| sdo:effectSize | sdo:isAssociatedWith | ods:Mean_Age |
| sdo:effectSize | sdo:collectedFrom | ods:EuropeanAncestry |

Each LOI has an *LOI question template* expressed similarly as the user question templates above. The LOI question pattern of each LOI is matched against the user question pattern through logical unification(Baader & Ghilardi, 2010). All LOIs that match are triggered and their methods are executed. The user’s question variables set up the LOI to find data and run computational workflows, as we describe in **Section 2.4**.

The question templates currently defined in NeuroDISK for investigating questions about the brain with neuroimaging genomics data in ENIGMA are:

- *What is the effect size of «Genotype» on «Region» «Brain Imaging Trait»?*
- *Is the effect size of «Genotype» on «Brain Imaging Trait» of «Region» associated with «Demographic Attribute»?*

We have also implemented the same questions with the addition of a filter, in this case filtering by specific ancestry labels of the cohorts:

- *What is the effect size of «Genotype» on «Brain Imaging Trait» of «Region» for cohorts of «Genetic Ancestry»?*
- *Is the effect size of «Genotype» on «Brain Imaging Trait» of «Region» associated with «Demographic Attribute» for cohorts of «Genetic Ancestry»?*

These question templates can be generalized further. For example we used genetic ancestry as a filtering criterion to replicate the discovery analysis of the original paper. However, the ontology can be easily extended to support other demographic characteristics or cohort level meta-data, for example, filtering by cohorts that use MRIs with a particular magnetic field strength or a particular genotyping chip.

Figure 5 illustrates how users specify their questions in the NeuroDISK user interface and the question pattern that results from it. Users do not need to be familiar with the ontologies or the structure of the data repository to pose their questions to NeuroDISK. Note that the genetic ancestry options are dynamically obtained from the data source and come from ODS-ENIGMA.

**Figure 5.**
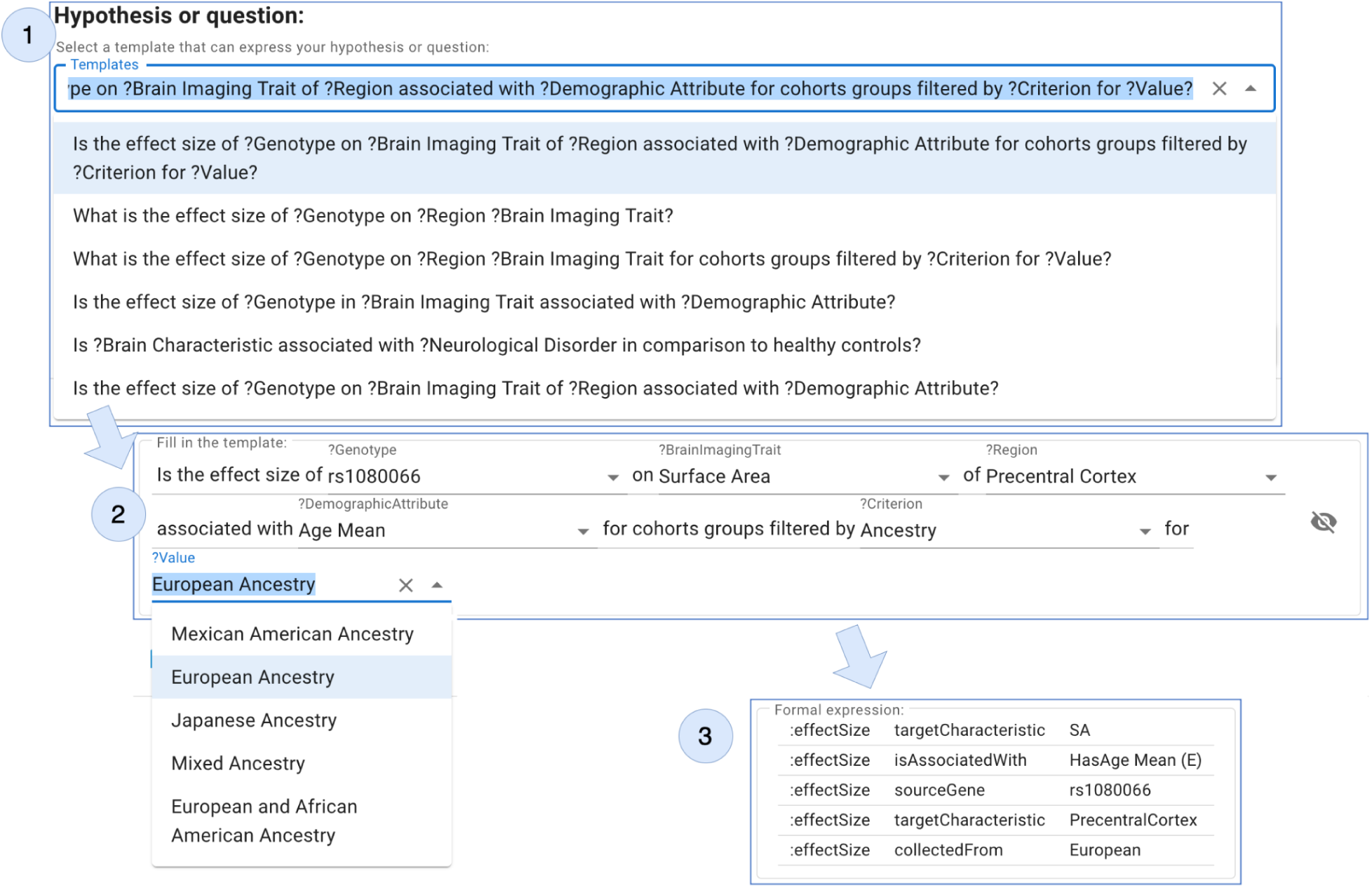
The NeuroDISK user interface for specifying questions. (1) A user chooses among a set of pre-defined question templates, which can be specific hypotheses with a posited outcome or simply exploratory questions. These templates contain variables and their possible instantiations. (2) The user chooses values for each of the question variables through pull-down menus. (3) The user’s question is turned into a question pattern consisting of RDF triples.

NeuroDISK can be extended with new question patterns and associated LOIs by users with advanced knowledge about the NeuroDISK framework. **Section 2.8** discusses different types of users and their interactions with NeuroDISK.

### 2.3 Organizing datasets through ontologies

Before we show how LOIs are triggered and executed, we describe in more detail how the ontologies are used to describe the datasets in the data sources queried by NeuroDISK.

The data should be described with semantic metadata that is rich enough to capture the terms that a scientist would use to identify what data they would consider relevant to a question or hypothesis. Many data sources have semantic annotations, using metadata vocabularies to describe characteristics of the data. In the case of NeuroDISK, the data source represents the available information from clinical research studies that collect data from a cohort of participants according to a specific study design with some inclusion/exclusion criteria, for example age range or clinical diagnosis. The inclusion criteria and other characteristics of the participants and the study design may be important for the user’s question. This description of the datasets is often obtained through semantic metadata, defined as part of one of several ontologies.

Figure 6 illustrates the main concepts in the ODS-ENIGMA ontology. It represents useful entities in the ENIGMA collaboration such as datasets, cohorts, organizations, protocols, instruments, software, working groups, projects, and persons, together with the relationships among them. It extends popular vocabularies for describing entities and actions (GuhaR, 2016) and the W3C semantic standard PROV(Gil et al., 2012) for provenance recording.

**Figure 6.**
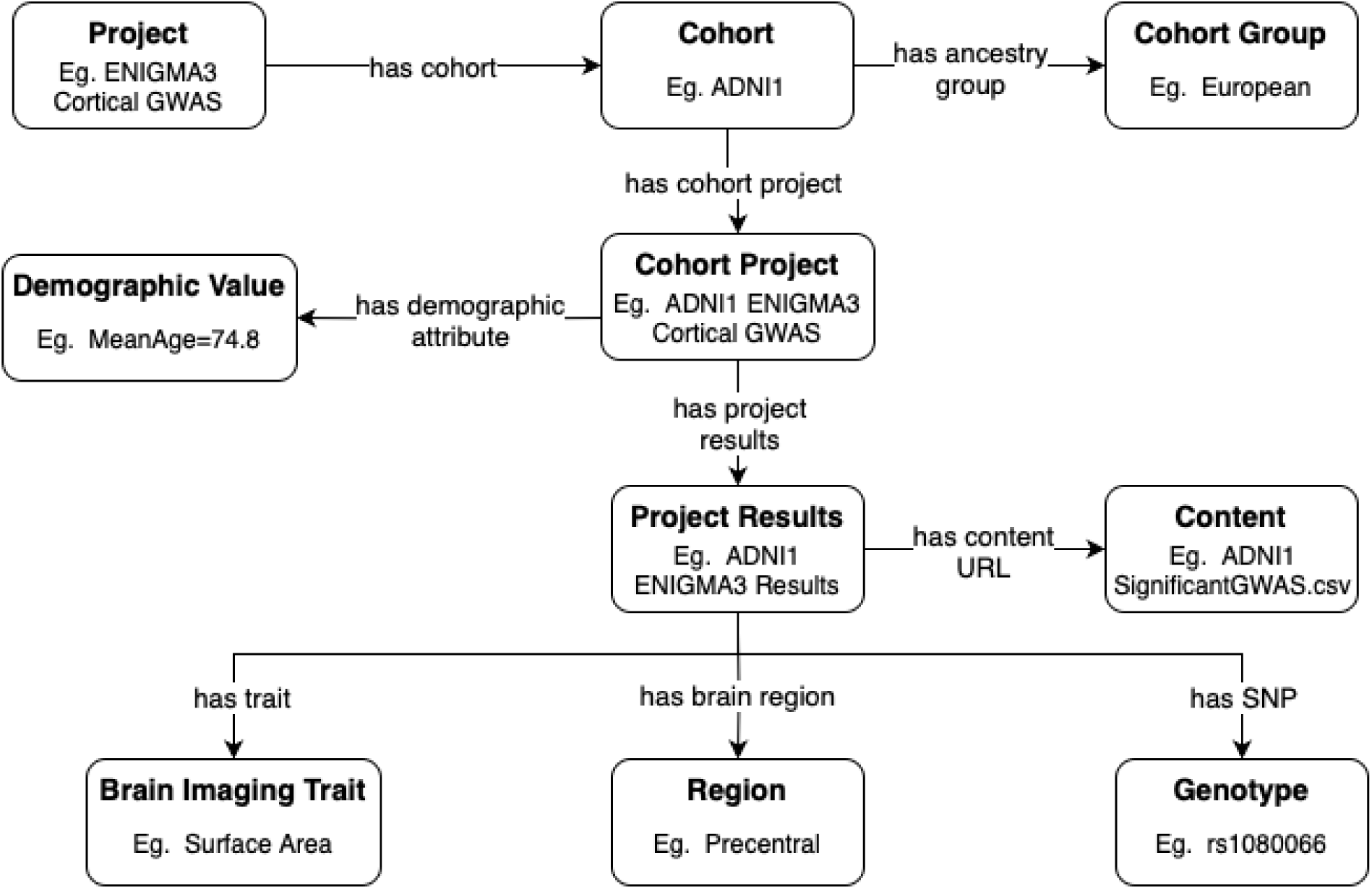
An overview of the main concepts in the NeuroDISK ODS-ENIGMA ontology (the DSO in NeuroDISK) that describe how cohorts and subsets of cohorts are used for specific projects.

In the ODS-ENIGMA ontology, a central concept is a *Working Group* that represents how several organizations in the ENIGMA consortium work together to analyze datasets on a particular topic of common interest. Each working group uses multiple *Cohorts* (defined as data from the set of participants who participated in a specific research study collected and maintained by a particular set of investigators); select data from these cohorts have been contributed by the working group members for collaborative analyses and projects, such as for the ENIGMA Cortical GWAS project.

Metadata about cohorts include the *Principal Investigator*, any *Covariates*, and *Brain Scan Data Type*, as well as relevant aspects of data collection, design type, and other statistical information such as the mean age of participants. Cohorts can also include subsets of the entire cohort to better define the data contributed to an individual project, or to better define different inclusion and exclusion criteria used in the same study (for different sets of participants).

Each working group organizes its members to collaborate in one or more *Projects*. A project uses data and information from multiple cohorts contributed by the collaborators. For example, the ENIGMA Cortical GWAS Project involves multiple cohorts and their associated data properties for the GWAS analysis of cortical measures. A cohort can be used in multiple projects and multiple working groups, but does not need to participate in all projects within any one group.

A project often uses the subset of a cohort that meets specific inclusion criteria, called a *Cohort Group*. A cohort can have multiple cohort groups, each with specific inclusion and/or exclusion criteria that defines them. For example, a cohort group may be the subset of a cohort considered “controls” or those without any neurological or psychiatric conditions. This distinction helps describe the different assessments that may be applied to particular subsets of the cohort; for example, “controls” may not have been asked to fill out questionnaires regarding medication use or have follow-up data, whereas the subset of individuals with a diagnosis of interest would have that information. A *Cohort Project Group* would then be considered the subset of the cohort group included for a particular project that meet all project inclusion criteria and pass quality control, such that exact statistics on included participants can be retained (e.g., N= 117, mean age=39.5). This can also be extended for different analyses within a project.

This representation for projects has the ability to capture provenance by maintaining descriptions of the cohort specifics that were used in a project at a specific point in time. This system allows for conserving cohort versions that were undertaken under certain cohort conditions. For example, the cohort ABCD can have a cohort project ABCD_proj_ENIGMA3_Cortical_GWAS, which contains all baseline cohort information available at the time of, and relevant to, the ENIGMA3 Cortical GWAS project. As more information is added to ABCD (such as new participants, covariates, assessments, etc.), this original Cohort Project remains untouched for its associated analyses. This makes documentation for past studies readily accessible.

To retrieve datasets described with this vocabulary, we express the queries using the W3C SPARQL semantic query standard.(Harris et al., 2013) For example, the following statements are used to create the query to retrieve all cohorts and their respective cohort projects and types of brain scans:

|  |  |  |
| --- | --- | --- |
| ?cohort | a | ods-e:Cohort |
| ?cohort | ods-e:HasCohortProject | ?cohortProject |
| ?cohort | ods-e:HasBrainScanDataType | ?ScanDataType |

The ODS-ENIGMA ontology is modular and composed of several smaller ontologies. **Table 1** gives an overview of these ontologies, which are documented online https://w3id.org/enigma.

**Table 1.** Overview of the current ODS-ENIGMA ontologies in NeuroDISK.

| Name | Description |
| --- | --- |
| Core Ontology | Main concepts of ODS-ENIGMA, elaborated further in other ontologies |
| Organization Ontology | Organizations (institutions) of investigators that contribute to ENIGMA |
| Cohort Ontology | Cohorts of study participants selected based on inclusion criteria |
| Demographic Ontology | Demographics for cohort datasets |
| Working Group Ontology | Dedicated working groups within the ENIGMA consortium |
| Project Ontology | Projects undertaken by ENIGMA working groups |
| Person Ontology | Researchers that participate in ENIGMA working groups and projects |

### 2.4 Finding data

Once a LOI is matched with a user’s question, the variables in the question are used to set up the LOI. The first step is to set up the data query to send to the data source in order to retrieve relevant datasets. Recall that the LOI has a data query template. To set up the LOI data query, LOIs have pre-defined *LOI variable mappings* between the LOI question template and the LOI data query template through their variables (Figure 7).

**Figure 7.**
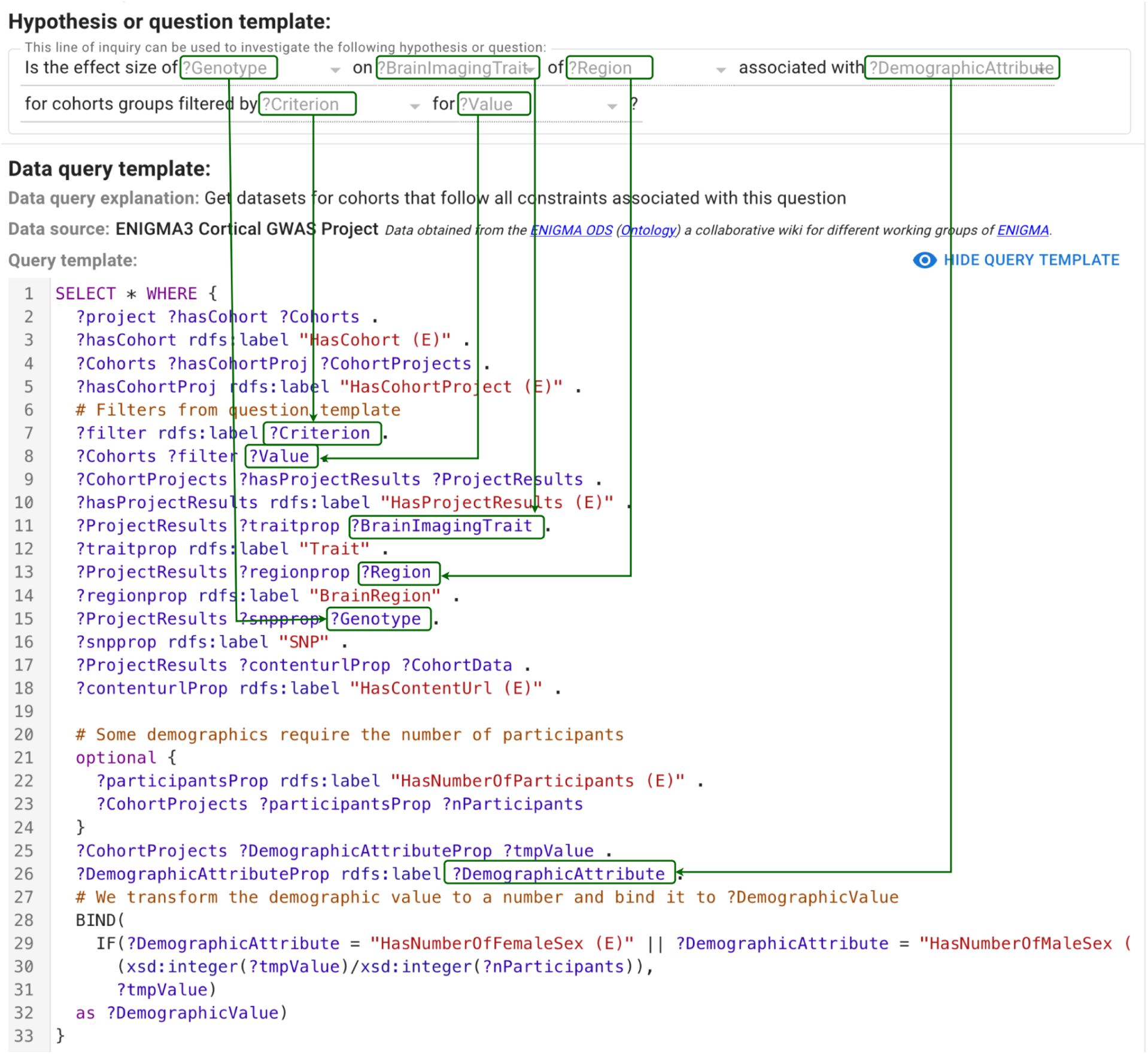
The LOI variable mappings between the variables in the question template and the variables in the LOI data query template indicate to NeuroDISK how to retrieve appropriate datasets from the data source. The data query template is a SPARQL query, and the actual data query will be completely specified once the user choices for the user question variables replace the corresponding LOI query template variables according to the variable mappings.

The LOI data query template has many variables that reflect how the data source is organized. In the case of ENIGMA, datasets are contributed by members who participate in ENIGMA working groups, where each member contributes data collected in their own clinical studies. Analogous to user questions and LOI questions, LOI data query templates have a *data query pattern* as a collection of triples that express the characteristics of the datasets sought.

The LOI variable mappings are used to incorporate the variable choices in the user question pattern into the LOI data query pattern to form the data query issued to the data source.

Figure 8 shows the data query in SPARQL for our running example, highlighting the LOI variable mappings that appeared in Figure 7.

**Figure 8.**
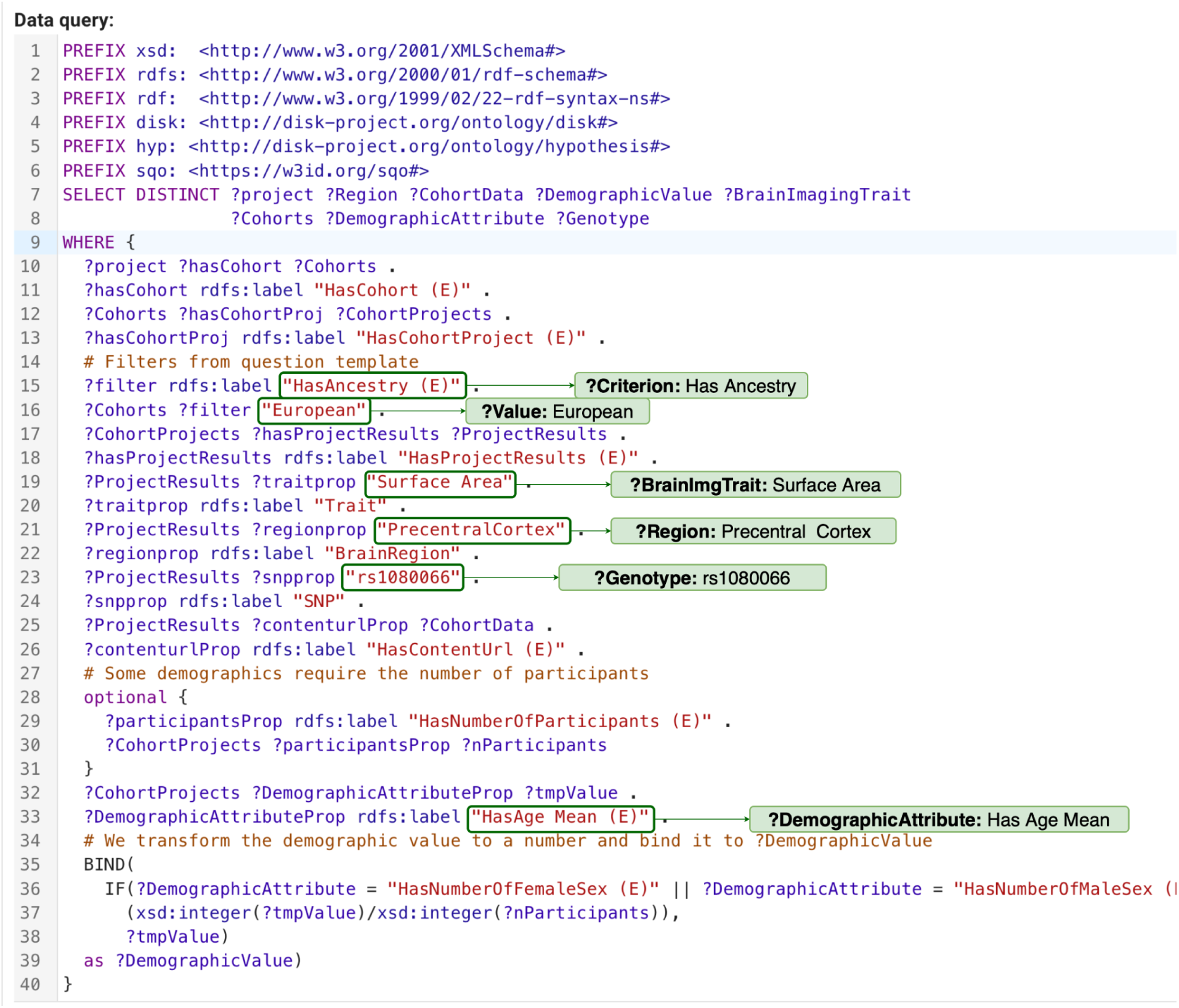
The data query generated from the user’s question, highlighting the relevant LOI variable mappings. The data query will be used to retrieve datasets from the data source that are relevant to a user’s question.

Once the data query is executed, the data source returns several datasets. The LOI indicates which variables in the LOI data query template would be useful for a user to see about those datasets. Figure 9 shows the results of the data query for the running example.

**Figure 9.**
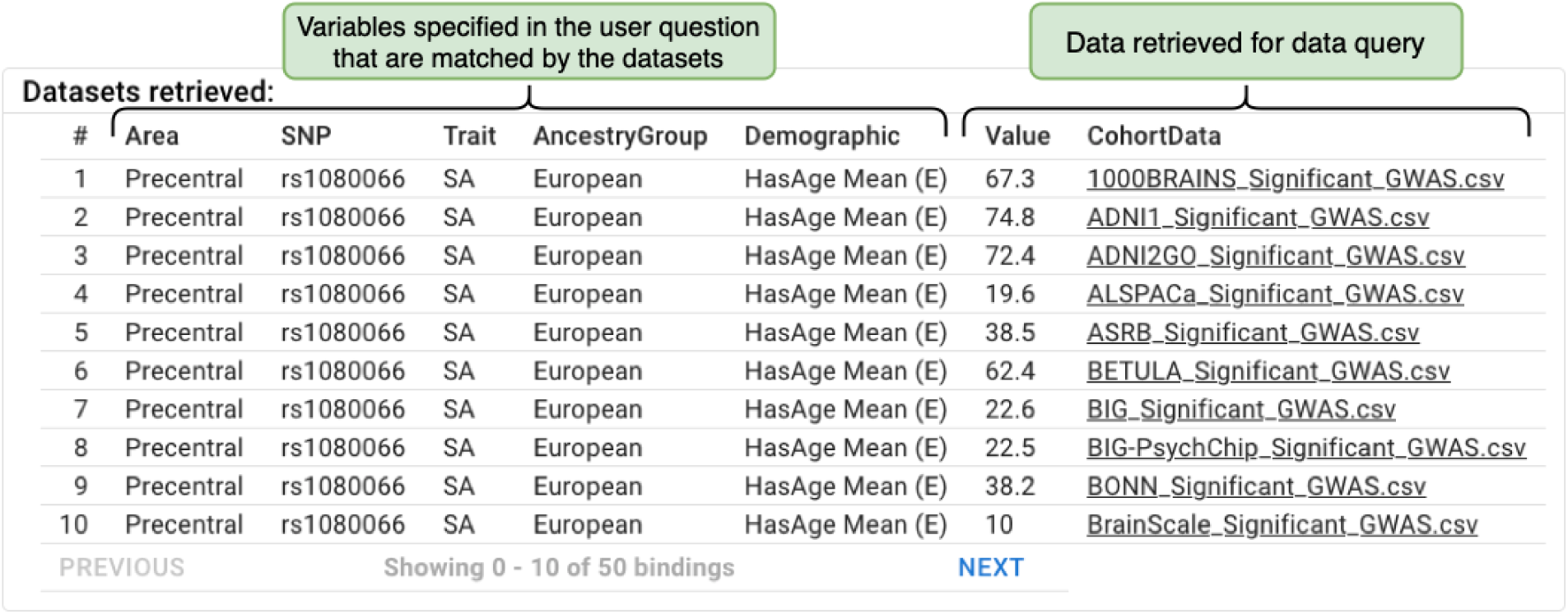
The cohort data that are retrieved, showing for each cohort the demographics and other information that appeared in the user’s question.

### 2.5 Analyzing data

NeuroDISK LOIs can analyze data in two stages: analysis and meta-analysis. Analysis typically refers to a method applied to a particular study (i.e, cohort). Meta-analysis typically refers to consolidating the results from individual analyses. This is done because the data within a study is often analyzed on site for privacy reasons. We represent them as workflows and meta-workflows respectively.

Workflows specify the steps and data dependencies needed to carry out a computational analysis, while meta-workflows capture the steps to aggregate the results from one or multiple workflows. When the execution of a workflow is completed, the provenance of all workflow execution results is captured, including the input and intermediate datasets as well as the software components used. Meta-workflows have access to all the outputs generated by the workflows, as well as to all the datasets retrieved by the LOI data query in case they are needed for the meta-analysis.

ENIGMA working groups often have projects that conduct meta-analysis, as individual subject information may not always be sharable. In this case, a statistical analysis such as a genome-wide association study (GWAS) is carried out locally at each of the sites that has stewardship over its own data, and only summary results are shared for meta-analysis. For our running example based on a project within the ENIGMA Genetics Working Group, NeuroDISK conducts the meta-analysis to combine the partial set of results of the individual site(cohort)-level analyses, and therefore, the currently implemented LOIs do not conduct any subject-wise analysis. In other words, the data source for the current examples includes a partial set of the results of the individual cohort analyses; when the data query retrieves these results, the LOI variable mappings pass the information to the meta-analysis workflow (Figure 10). The results of running NeuroDISK for a user question are highlighted in Figure 11, explaining the LOI used, the datasets retrieved, and the results obtained. These explanations are generated from the provenance records that DISK keeps, including provenance records for the workflow executions that the workflow execution system provides as described below.

**Figure 10.**
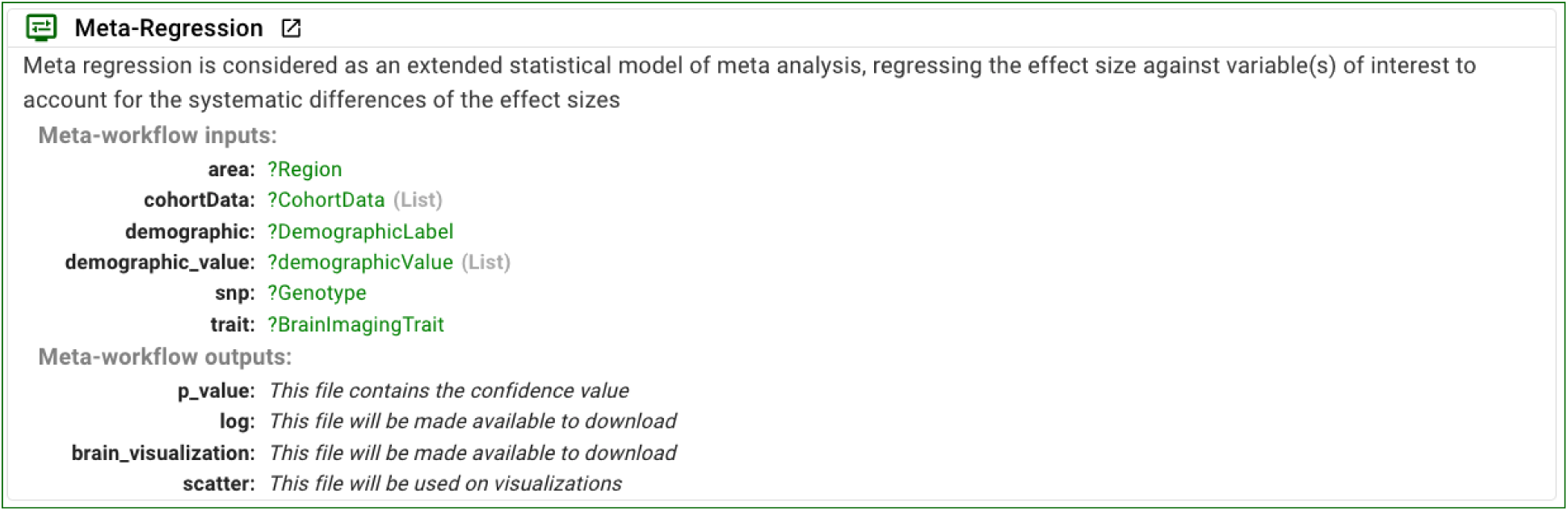
The LOI’s meta-analysis is done through a meta-workflow, in this case for meta-regression. The LOI variable mappings are used to take the results of the data query and set the inputs to the meta-workflow.

**Figure 11.**
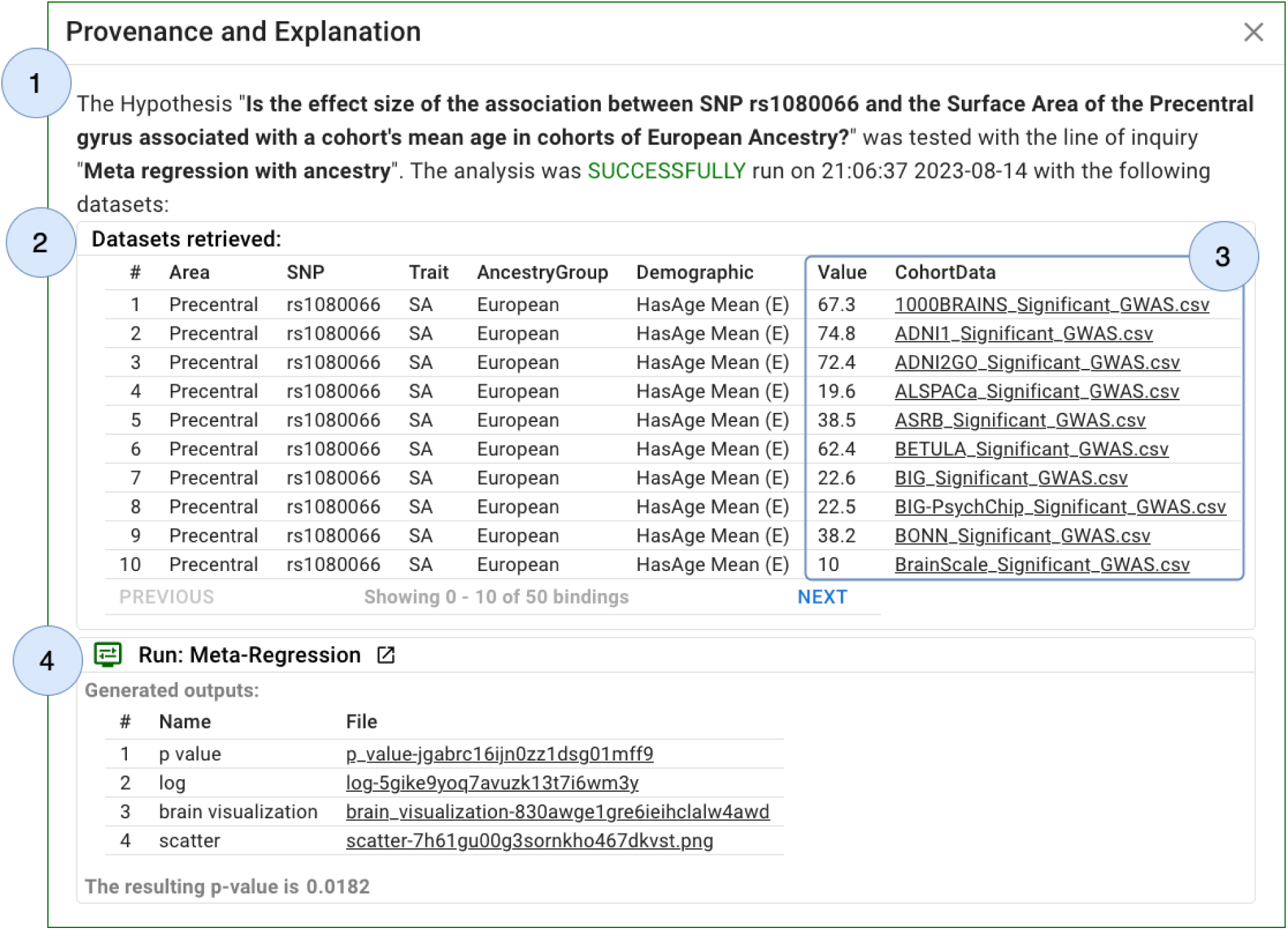
NeuroDISK captures the provenance of all execution results of the analysis and meta-analysis workflows and uses it to generate explanations for users. In this case: (1) the original question or hypothesis is shown along with the name of the selected LOI; (2) the datasets retrieved corresponding to several cohorts; (3) datasets and dataset characteristics that were input to the meta-analysis; (4) the results of the meta-analysis, which include a confidence value as well as visualizations of the results.

### 2.6 Continuously updating results

NeuroDISK continuously checks for new information that could be used to answer an active query; new information may be new cohort data added to the data source or new workflows to reflect new analysis methods. In such cases, NeuroDISK re-runs the LOI to update the findings. Therefore, there may be several runs of the same LOI in response to a standing user query, all of which are captured and saved.

Figure 12 illustrates how users see this process. In this case, initially 10 datasets are available, and eventually up to 50 cohorts’ datasets become available in the data source.

**Figure 12.**
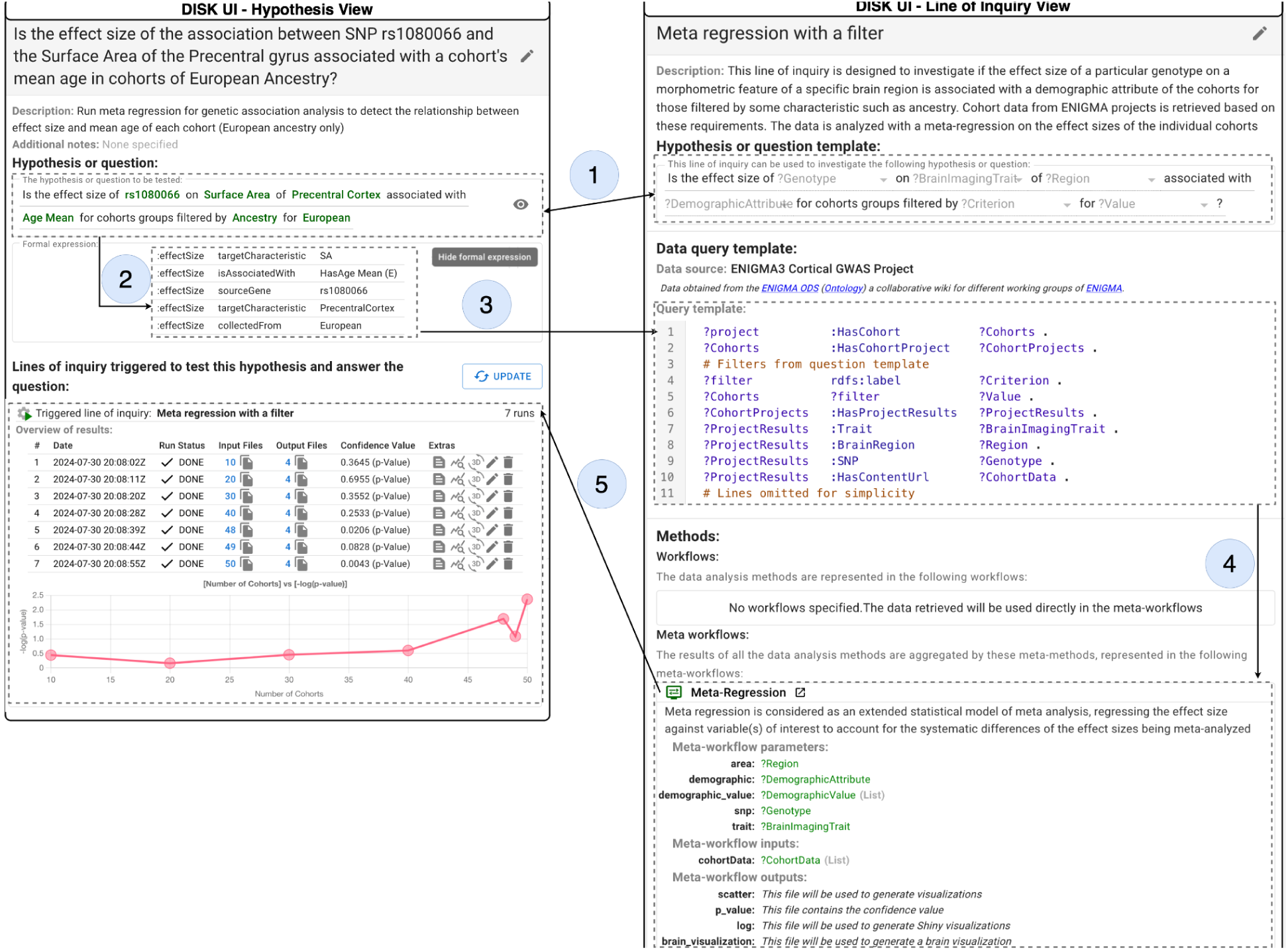
NeuroDISK continuously checks if the results for a user question have changed due to new data becoming available or due to workflow or meta-workflow updates. (1) The standing user question continuously triggers the matched LOI; (2) the user hypothesis contains variables; (3) those variables are used to set up a query to retrieve relevant data; (4) the data is then analyzed with workflows; and (5) the LOI is re-executed every time that additional data is retrieved from the data source, and the user can see how the results change. On the left is the user’s view of the hypothesis and the results over time. In this case we show the −log10 of the p-value against the iterations, and overall highlight smaller p-values or an increase in confidence (higher points on the plot) with added data points. Users can ask for details on each run, and can request to see the LOI as shown on the right.

### 2.7 Implementation of NeuroDISK

An integrated overview of the components of NeuroDISK is shown in Figure 13. We distinguish among *users* who formulate questions and receive the results, *advanced users* who define the types of questions and corresponding lines of inquiry, and *developers* who integrate new data sources and workflow software into the framework.

**Figure 13.**
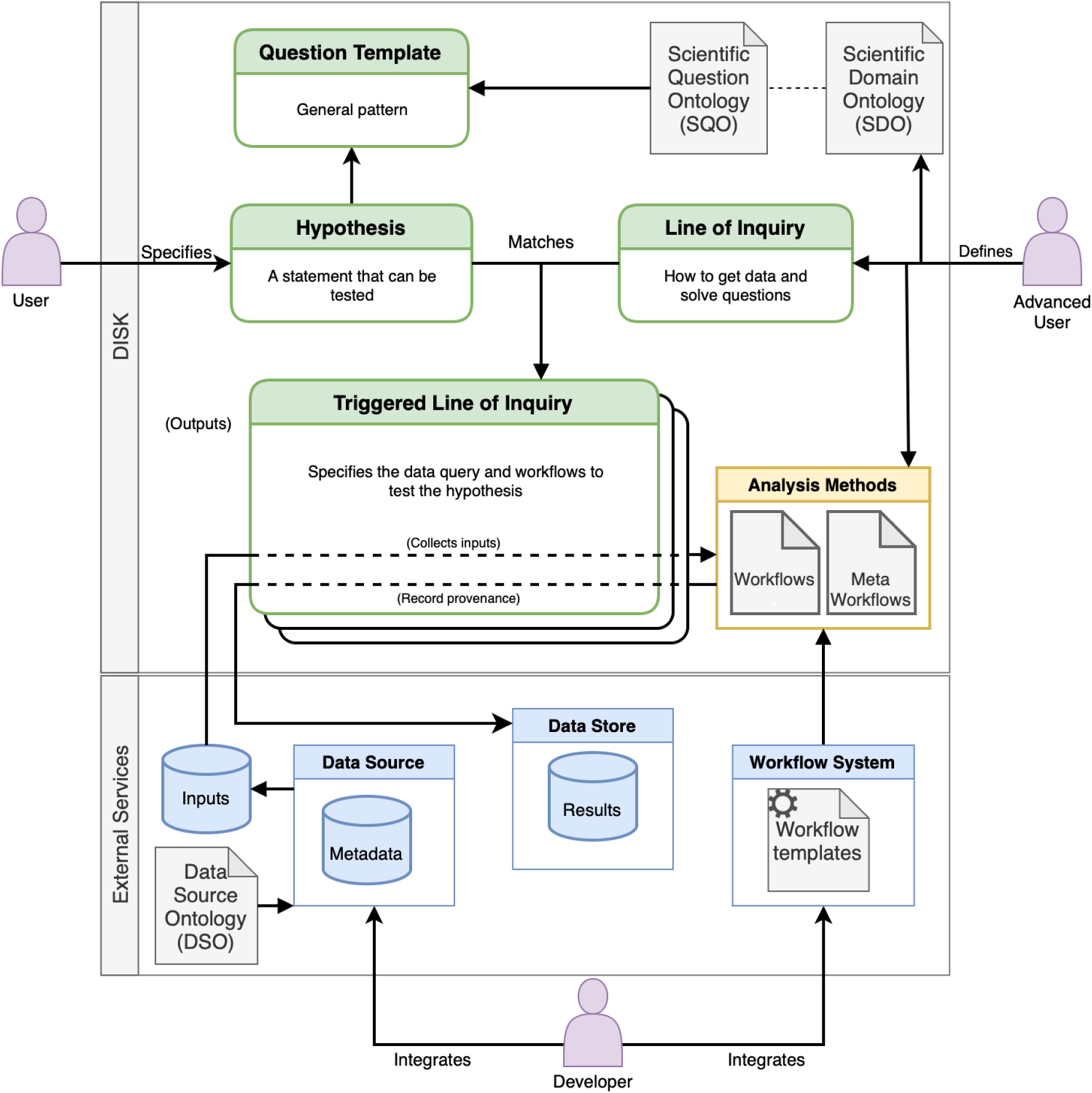
A diagram of the architecture components of DISK. Users specify hypotheses or questions that conform to pre-defined templates. Advanced users have added pre-defined LOIs that are triggered when they match the user’s question. The LOIs generate queries to retrieve data from a data source, and the data is analyzed through workflows and meta-workflows. A developer can set up new data sources and new workflows in the DISK back-end.

In NeuroDISK, the data source is implemented in the Organic Data Science (ODS) framework(Gil et al., 2015), an extension of Semantic MediaWiki(Krötzsch et al., 2007). This enables users to view metadata of different datasets and to add new metadata as needed. In NeuroDISK, we set up a separate ODS site per ENIGMA working group, since each group operates largely independently, although these largely share the same ontologies. The ODS data source we make available as part of this publication is one which contains published information(Grasby et al., 2020) from the cortical GWAS project of the Genomics Working Group.

In NeuroDISK, workflows and meta workflows are executed through WINGS(Gil et al., Jan.-Feb 2011, 2011), an intelligent workflow system that propagates semantic constraints to ensure that the analysis is valid for the data at hand, and to set up parameter values that are appropriate for the input datasets. WINGS exports provenance information to DISK, so that it is accessible to its users.

The architecture of DISK is modular, and other data sources and workflow systems can be integrated. Figure 14 shows the data adapters and APIs defined for DISK.

**Figure 14.**
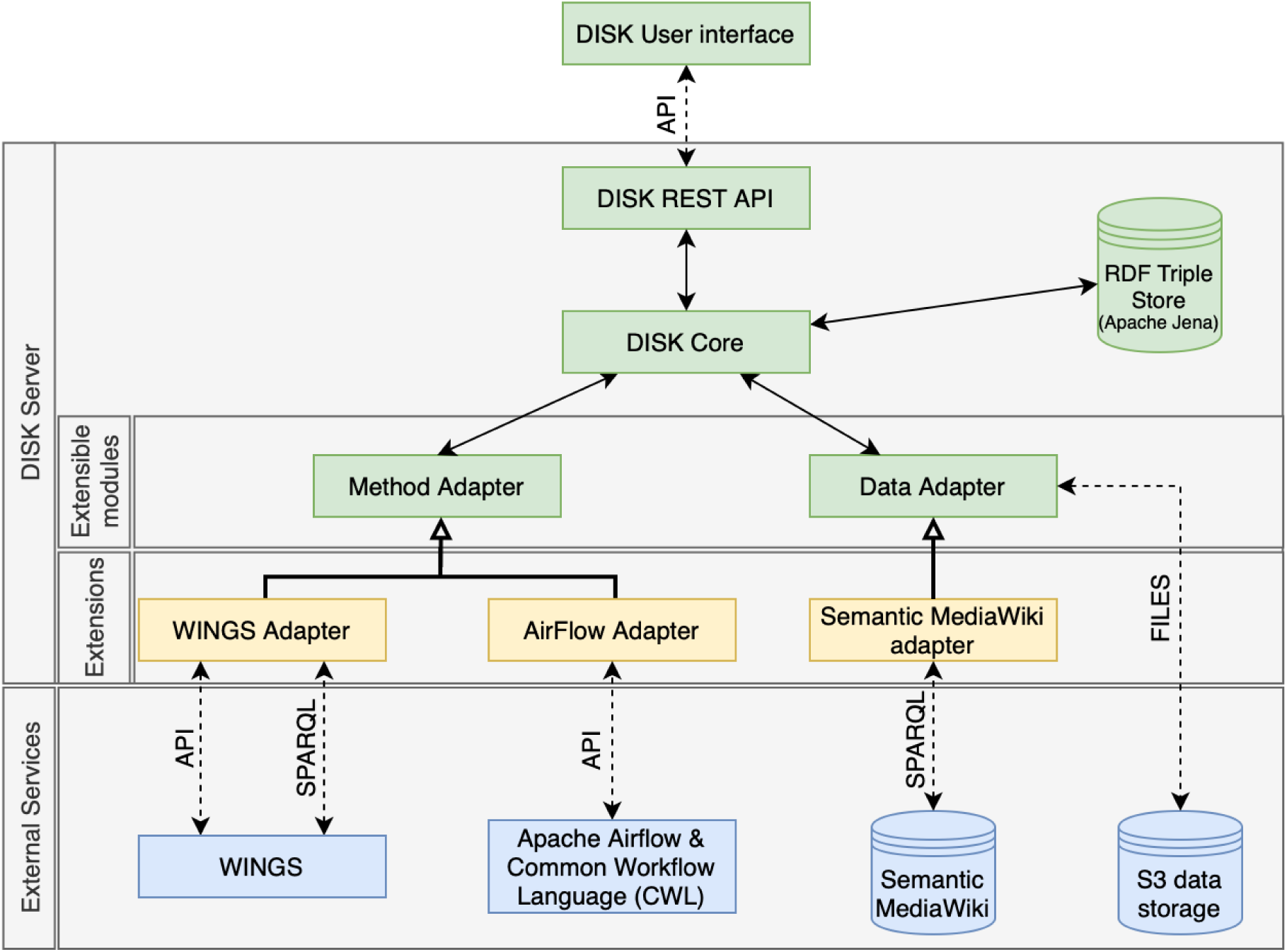
A diagram of the DISK APIs and adapters that enable interoperability with other data sources and workflow systems. DISK provides abstract classes to implement both data adapters (for data sources) and method adapters (for workflow systems). The current implementation of DISK includes method adapters for two workflow systems, WINGS and AirFlow, and one data adapter for the Semantic MediaWiki platform.

NeuroDISK is fully implemented, and links to a web portal, system software, ontologies, and other materials are included as Supplementary Materials. The system can be installed as a software container or building from source code. The system is documented in detail with guides for users, developers, and system administrators.

## 3. Results

We demonstrate the use of NeuroDISK to simulate hypothesis-driven discovery and continuous result updates using and extending a well-known ENIGMA publication(Grasby et al., 2020), in which researchers from institutions around the world participated in this study by following specific standardized protocols to: 1) extract neuroimaging derived traits from the cortical brain surface; 2) impute the genotyping data 3) quality control all the data; and 4) run a series of linear association models (or mixed effects models depending on population structure) to identify genome-wide associations with cortical brain imaging measures. 70 total brain imaging traits were assessed genome-wide (∼18,000,000 genetic variants). The results from cohorts of European ancestry were meta-analyzed together using an inverse-variance based weighting. Replication analysis was performed by meta-analyzing results from the ENIGMA cohorts with that of a single large cohort, UK Biobank, with over 10,000 individuals. These final meta-analyzed results were then used to demonstrate generalizability in other cohorts of individuals from non-European ancestry.

While final meta-analyzed results are available for the full genome-wide and image-wide set of variables on the ENIGMA website, it has been suggested that raw cohort level results can be used to identify an individual participant(Cai et al., 2015; Homer et al., 2008). Therefore, here we work only with a subset of results. The subset of summary statistics includes the effect size, standard error, and p-value calculated for 14 of the significant associations discussed in the paper between specific single nucleotide polymorphisms (SNP) and surface area or thickness for cortical regions of interest. The individual cohort-level results were made available as part of the Forest plots in the supplementary materials of the original publication. The set of results per cohort were uploaded into NeuroDISK as a single file, with the results of each SNP as an individual row, allowing the set of findings made available to easily be extended in the future. While all 14 associations are available for users to peruse, our running example uses the most significant association identified in the original publication – the SNP rs1080066 and the surface area of the precentral cortex. We also capture meta-data related to the sample size, sex distribution, mean age, and ancestry information available from participating cohorts.

We first reproduce the results of the Grasby et al(Grasby et al., 2020) publication with a meta-analysis of all effect-sizes as weighted by the inverse variance of the effects. Next, we show how cohort-specific meta-data can be integrated to ask a novel question of the data, which may help identify discrepancies in results across cohorts. Finally, we add data from the Adolescent Brain Cognitive Development Study (https://nda.nih.gov/abcd)(Jernigan et al., 2018) a cohort that was not available as part of the original 2020 paper, and highlight how incrementally adding data can change results, altering confidence in the original meta-analysis and the novel meta-data based analyses. We detail these results below.

### 3.1 Replicating the published ENIGMA GWAS meta-analysis

In Grasby et al.,(Grasby et al., 2020) we and over 300 co-authors performed a standardized genome-wide association study meta analysis as described above. Here, we show that when the individual level cohort summary results for the SNP of interest are stored in the ODS database, we can query the association with NeuroDISK and trigger a meta-analysis workflow. The query specifically searches for association results of SNP rs10810066 on the precentral surface area, filters for cohorts of European ancestry, and performs a meta-analysis. Our meta-analysis workflow also generates a Forest plot for visual comparison of effect sizes per cohort. Users are presented with the individual datasets that the query returns and have the option to remove individual cohorts. Here we show that a meta-analysis of all discovery ENIGMA cohorts and the UK Biobank results yields results very similar to that of the published work (Figure 15). The slight discrepancy is due to the fact that in the publication the ENIGMA cohorts were meta analyzed together first, then UK Biobank was meta-analyzed with those results, where as here we meta-analyze results from all cohorts (including UKBiobank) together. We show this by comparing the results of two associations (rs10810066 on the precentral surface area) from both the original publication and that of NeuroDISK’s reassessment with and without UK Biobank (**Table 2**). We show that the resultant effects are nearly identical without UK Biobank, but differ slightly in significance with UK Biobank due to the difference in how the cohort was included, as the Grasby et al paper meta-analyzed the result of all 48 ENIGMA cohorts (considered discovery) with that of UK Biobank (considered replication), while here we meta-analyze all 49 cohorts simultaneously.

**Figure 15.**
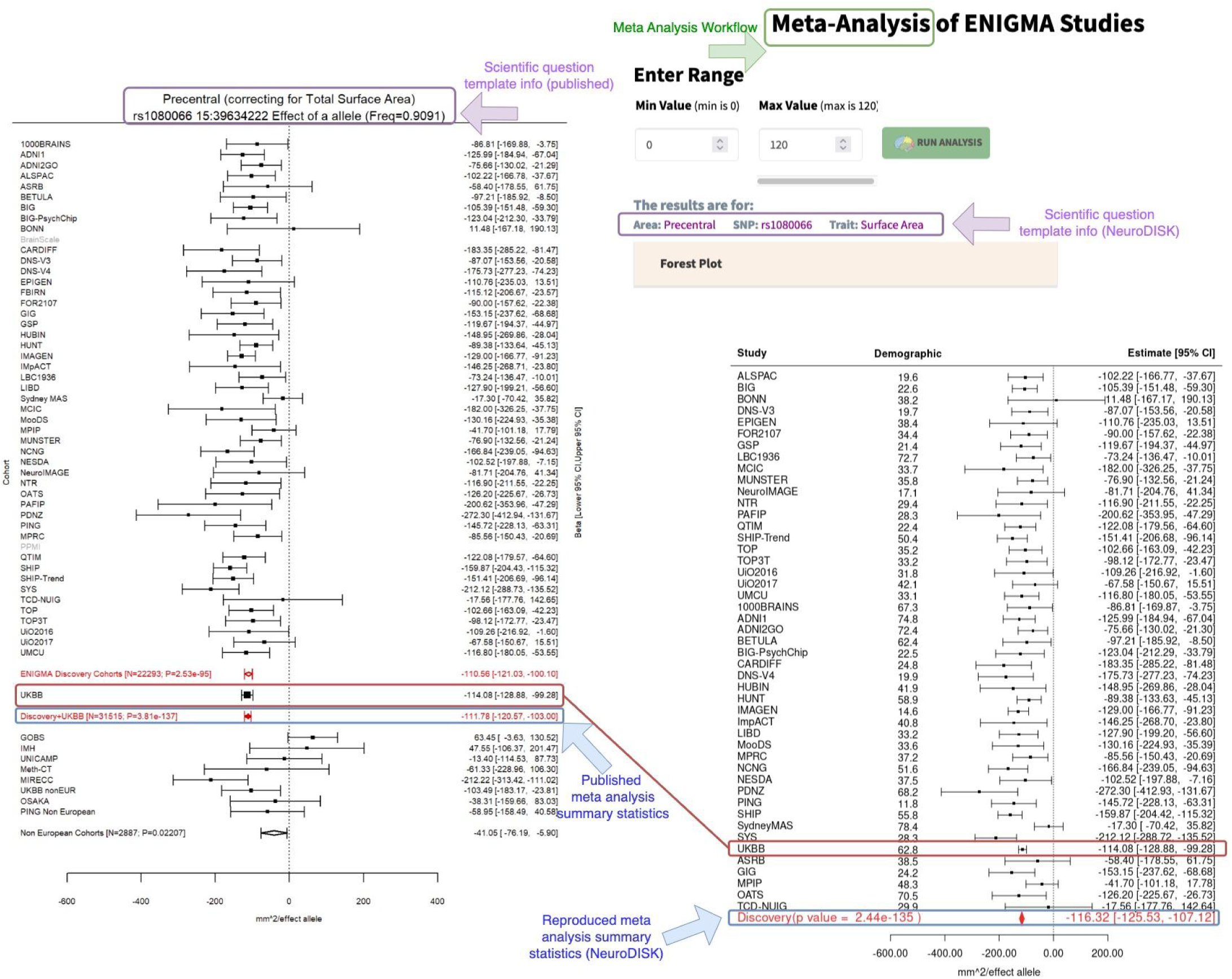
Meta analysis workflow results to validate the NeuroDISK framework can reproduce the published meta analysis results(Grasby et al., 2020). Left: Published Forest plot for the effect size of SNP loci rs1080066 on the precentral surface area and from supplements of published results(Grasby et al., 2020). Right: Forest plot using the NeuroDISK meta analysis workflow highlights successful replication.

**Table 2.** Reproduced genetic association results for precentral surface area and rs1080066 for ENIGMA discovery cohorts with and without UK Biobank.

| Precentral surface area and rs1080066-A allele | ENIGMA Discovery Cohorts |  | ENIGMA Discovery + UK Biobank |  |
| --- | --- | --- | --- | --- |
|  | Published | Replicated | Published | Replicated |
| Effect size (unstandardized beta) | -110.56 | -110.51 | -111.78 | -116.32 |
| P value | $2.53 \times 10^{-95}$ | $6.70 \times 10^{-95}$ | $3.81 \times 10^{-137}$ | $2.44 \times 10^{-135}$ |

### 3.2 Asking and answering novel questions

When working across such diverse cohorts and performing meta-analyses, researchers may be interested to know whether there are aspects of particular datasets that are driving the associations. For example, is the effect of a genetic association a function of the age of a specific cohort? We can ask and answer such questions within the NeuroDISK framework. Specific association results and covariates are retrieved from the ODS, and meta analysis or meta regression workflows are provided as options. In the case of rs1080066 and precentral surface area, the meta regression workflow corresponds to ‘Is the effect size of *rs1080066* on *Precentral SA* associated with *mean age of the cohort*?’ We performed a meta-regression to determine whether there was an association between the effect size of the SNP’s association with the cortical structure and mean age of each cohort. Our meta-regression workflow weighs the effects of the cohort by its sample size and regresses the effects against the cohort-specific factor, in this case for mean age. The workflow includes a scatterplot as a visual output and allows the user the option to filter cohorts by limits on the factor. For example, in Figure 16 we show results of the meta-regression across the full set of cohorts (left) and after filtering to only include cohorts with mean age under 60 years (right). Although not shown, filtering can also be done as part of the meta-analysis in **section 3.1**.

**Figure 16.**
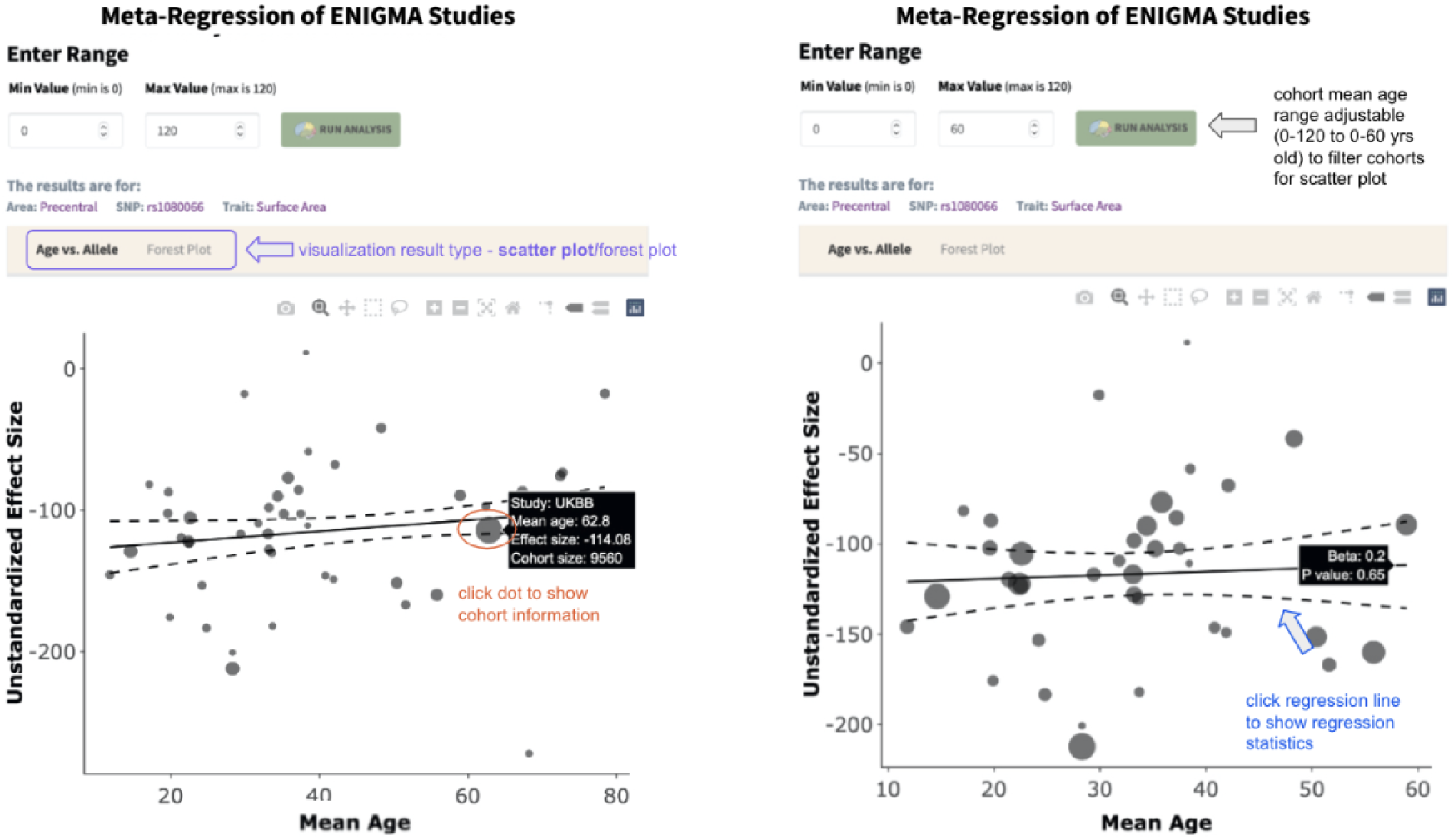
Meta regression workflow results show a scatter plot displaying the association between the effect size of an association of interest, here being the effect of SNP rs1080066 on precentral surface area (y-axis), against the mean age of each cohort (x-axis). The visualization is based on R Shiny (https://www.rstudio.com/products/shiny/) and is interactive such that users can click any point to check the cohort information. Clicking the regression line shows the beta value and p value of association. The size of each point is a reflection of the sample size of the cohort. (Right) Users can also adjust the demographic variable of interest to selectively display cohorts that fall within a specified range, ie., the mean age between 0-60 years old.

### 3.3 Continuous updates of results

The NeuroDISK framework can re-execute queries as more data becomes available. We demonstrate this re-execution through continuous retrieval of data and updating of results by artificially simulating the availability of new data over time. We subsampled the available data, starting with 10 cohorts (N = 10) and then iteratively adding 10 more cohorts at a time (N = 20, 30, 40, 48) until we reached 48 total cohorts, equal to the total number of cohorts in the original ENIGMA discovery cohort (not including UK Biobank). We then added UK Biobank, which was analyzed separately in the original paper (N = 49). Lastly, we added a new external cohort, the Adolescent Behavioral Cognitive Development (ABCD) dataset, which was not in the original publication, so associations were calculated separately and uploaded onto the ODS (N = 50 cohorts).

ABCD is the largest longitudinal neuroimaging study for child health in the United States, which has recruited more than 10,000 children around 9 to 10 years old(Casey et al., 2018) with deep genotyping using the Affymetrix Axiom Smokescreen Array. We extracted demographic information such as age, sex, and scanner information (SIEMENS/GE/Philips) for the baseline (N = 11,362) data. T1 weighted MRI from ABCD 4.0 release data were processed by the ABCD group(Hagler et al., 2019) and baseline cortical surface area or thickness measures were extracted using Freesurfer version 7.1.1 available in release 4.0. We calculated the first four components of MDS using the ENIGMA protocol and followed the ENIGMA genetic imputation protocol and QC criteria (Minor Allele Frequency < 0.01; Genotype Call Rate < 95%; Hardy-Weinberg Equilibrium < 1×10^-6^). We filtered for individuals of European ancestry as in the ENIGMA Genetics Imputation protocol available online (https://github.com/ENIGMA-git/Genetics/tree/main/ENIGMA2/Imputation), where a radius of 0.0066 was set around the first, second and third MDS components of the CEU centroid, resulting in a sample size of N = 5,202. The distance was calculated by covering 95% of the first, second and third MDS components of the European centroid data. We used GCTA(Yang et al., 2011) to estimate the genetic relationship matrix (GRM) from 452,544 SNPs, which was subsequently used in a linear mixed model regression of the specific SNPs of interest on the brain regions of interest using GCTA-MLMA(Yang et al., 2014) and including age, sex, and scanner manufacturer (Siemens, GE, or Philips) as fixed covariates along with the first four genetic components derived from multidimensionality scaling, as performed in the original ENIGMA publication(Grasby et al., 2020).

As ABCD was not used in the original publication, these genetic association tests were run separately before corresponding genotype-phenotype association results were uploaded onto the ODS for follow-up continuous meta-analysis and meta-regressions.

Figure 17 shows the meta-regression results of the consecutive runs with increasing sample sizes and shows the trend of the outputs mapped against the number of data inputs. On the UI itself, users can see the final p-value of the association and access the plots (Figure 16) to help visualize the individual data points. As the number of cohorts increase to 40 or more, the effect of mean age on the effect of the SNP on the cortex appears to reach nominal significance (p=0.02), showing a trend towards younger cohorts having a greater genetic effect. We do note that no stable trend is being observed, and more data will be needed to establish greater confidence in the significance of the association.

**Figure 17.**
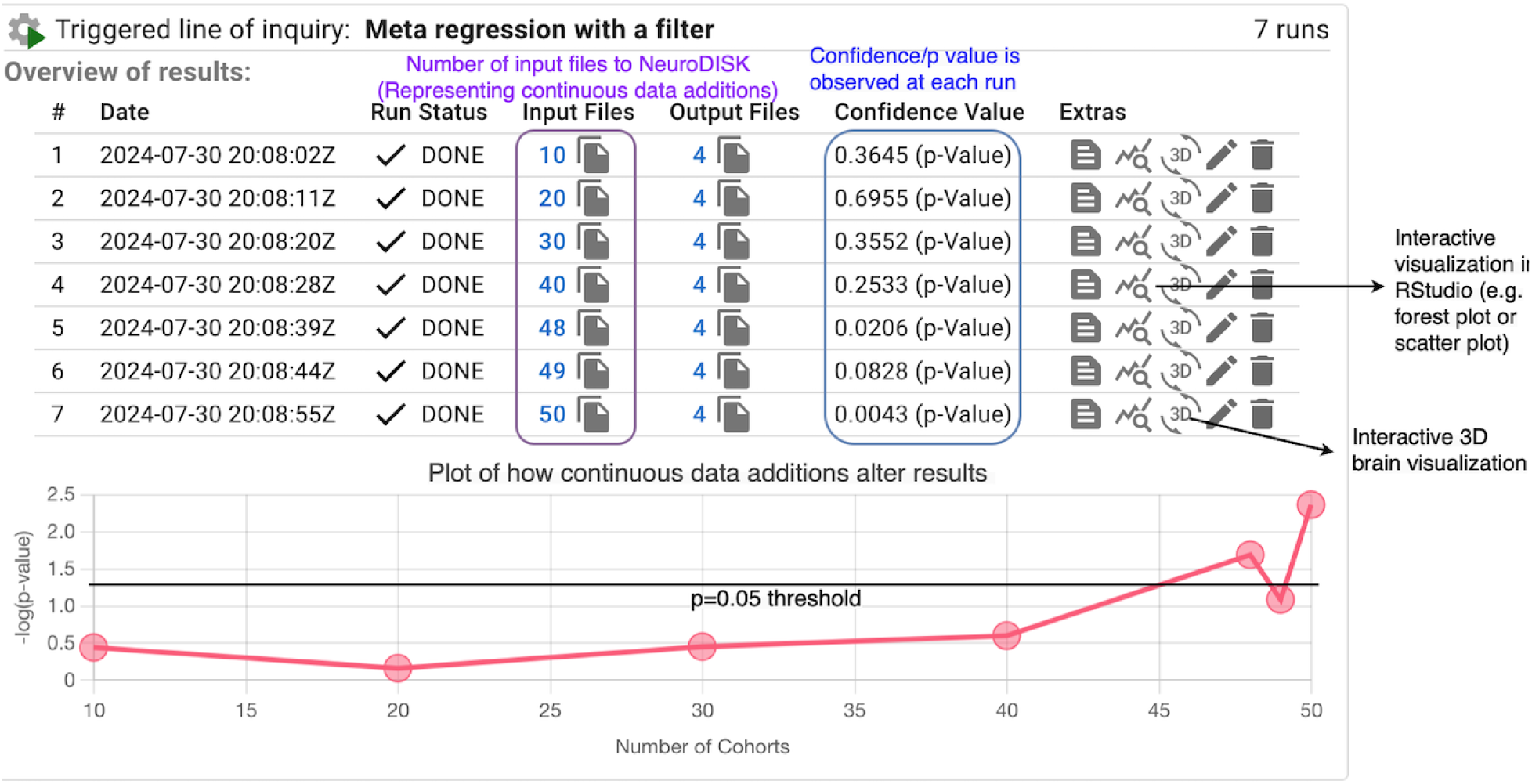
Continuously updated findings. The NeuroDISK user interface shows results across all runs in one location for ease of comparison. We display the meta-regression results of continuous runs of the effect of SNP rs1080066 on the precentral surface area versus mean age of the cohort. To demonstrate a continuously growing database of cohort data, the first 10, 20, 30, and 40 samples were taken at random from the initial list of 48 cohorts of European ancestry involved in the original ENIGMA cortical GWAS publication(Grasby et al., 2020). The 49th cohort was the UK Biobank dataset which served as a meta-analysis replication in the original publication, and the 50th included cohort was ABCD, a cohort not involved in the original publication. We notice that as the number of cohorts increase, the effect of mean age on the effect of the SNP appears to reach nominal significance, although as the result hovers around the nominal p=0.05 threshold, more data will be needed to ensure stability of the effect. The −log(10) of the p-value is plotted on the y-axis, so points higher along the y-axis are more significant.

In summary, our results show that NeuroDISK can perform data analysis continuously and automatically, and our methods yield results that replicate the original publication and allow users to delve deeper with the data and ask additional questions of summary results that were not answered in the original study. NeuroDISK is ideal for meta-analytical studies, as demonstrated in this work, but can also be extended to large-scale coordinated analyses using individual subject level data points.

## 4. Discussion and Conclusion

We have presented a novel AI approach to scientific problem solving that captures how scientists pursue inquiry-driven discoveries through explicit knowledge structures called lines of inquiry. We use AI techniques to represent how scientists pose questions or hypotheses, how they characterize the data needed to address a given question, and what methods are appropriate to analyze the data found. We use AI reasoners to execute expression matching, variable assignments, constraint propagation, and other forms of problem solving needed to set up and run lines of inquiry. Our approach is implemented in NeuroDISK, which extends the general-purpose DISK framework with lines of inquiry for multi-site imaging genetics, including datasets with descriptive metadata and workflows for analysis. We demonstrated NeuroDISK by reproducing the results of a well-known publication following our approach, and showed how the results can be updated automatically when new data becomes available.

NeuroDISK has demonstrated important capabilities for the automation of inquiry-driven discovery, specifically:

- *Guiding scientists to specify neuroimaging genetics inquiries that can be tested with available datasets*. NeuroDISK can guide scientists to specify an inquiry (question or hypothesis) by selecting from possible questions that can be answered using available datasets. This is possible because NeuroDISK uses question patterns that are tied to ontologies representing the kinds of terms in questions that could be answered with the data available. The underlying DISK framework provides a general ontology of scientific questions and hypotheses that provides overarching concepts that guided the design of the neuroimaging genetics question ontology in NeuroDISK.
- *Automatically selecting among possible approaches that represent how to answer common types of neuroimaging genetics inquiries with existing datasets*. NeuroDISK contains lines of inquiry that express the general approaches that a scientist would pursue for a given inquiry, including what kinds of datasets to use and how to analyze them. NeuroDISK lines of inquiry provide the basis for taking a scientific question and automatically generating a query to a data repository, then setting up and running analyses for the data found, and finally consolidating the results to answer the question.
- *Automatically finding data by generating data queries that describe characteristics of datasets from existing neuroimaging cohorts that make them useful for different types of scientific questions*. NeuroDISK includes ontologies to describe cohorts datasets collected in neuroimaging studies and to describe subsets of those cohorts that have been used in ENIGMA projects. Dataset characteristics include genotypic, phenotypic, and other demographic information. These representations enable NeuroDISK to automatically retrieve and reuse available datasets. This approach is consistent with the FAIR data principles(Wilkinson et al., 2016) by providing a descriptive metadata to find, access, integrate, and reuse datasets. For NeuroDISK, metadata about project cohorts is in the data repository (the ODS Wiki), in a machine readable format and accessible through APIs.
- *Automatically apply methods by elaborating and executing appropriate steps to analyze the available data to investigate an inquiry posed by users.* NeuroDISK reasons the representations of scientific questions and the representations of lines of inquiry in order to automate this process. In this way, NeuroDISK generates data queries to retrieve appropriate datasets and set up executable workflows for analyzing that data with the required inputs. This may be decomposed into analysis, done with workflows, and meta-analysis done with meta-workflows.
- *Updating results by continuously monitoring the available data sources and triggering new runs on the relevant lines of inquiry*. NeuroDISK considers standing user inquiries to reconsider lines of inquiry that query data sources for new datasets. When new data is returned, the analyses are executed again and the results are updated.
- *Presenting the findings through explanations and summaries of the analyses conducted*. NeuroDISK generates explanations of how a result was generated based on the provenance records of the execution of a triggered line of inquiry and its associated queries and workflows. When new datasets are added, NeuroDISK summarizes the differences among all the triggered lines of inquiry so that the changes in findings can be tracked.

Our system has certain limitations given that NeuroDISK was designed with a specific scope based on the select ENIGMA projects that we targeted. We designed NeuroDISK with modularity in mind so that it can be easily extended for other datasets, projects, and studies. For example, new ontologies and workflows would need to be created for other neuroimaging data of different modalities (diffusional MRI, functional MRI etc.).

NeuroDISK is designed to support scientists to explore questions and hypotheses. Rather than executing the above steps and processes manually, scientists can rely on NeuroDISK to automate the process and present the results in context. This may save scientists significant time and effort. It also has assurances that the analysis is valid, since it uses proven methods as represented in NeuroDISK. In addition, NeuroDISK reconsiders a scientist’s question whenever new relevant data becomes available. This would add a novel dimension to scientific research, namely obtaining updates of published findings as more data is collected in various neuroimaging studies over time. This could help increase the confidence of published findings or lead to revisiting previously explored questions to refine the scope of a past investigation. As new datasets are incorporated into the analysis, a scientist can consider more specific questions such as filtering by a demographic characteristic where enough data is now available. Finally, NeuroDISK could be extended to automatically generate visualizations that highlight the results in different populations so that scientists can consider potential future investigations.

Beyond individual users, there are many benefits of a framework like NeuroDISK for use in scientific collaborations such as the ENIGMA consortium. First, method validity: lines of inquiry and workflows in NeuroDISK would be used for many analyses and reviewed by multiple users. This would be in contrast with current practices where individual researchers establish their own methods and software. Second, dissemination of new methods and method reuse: Our system makes it easy to run existing methods as there would be no prior learning needed to apply them. Third, transparency and reproducibility of the analyses: Everyone can access the provenance traces for any analyses in the collaboration. Fourth, promoting FAIR data and FAIR method practices: AI reasoners need explicit metadata and representations that are also useful for human scientists. Fifth, comparison of experimental results would be facilitated as the provenance and lines of inquiry would be comparable side by side.

Adopting a framework such as NeuroDISK could raise concerns about stifling scientific creativity, since the framework would run the same workflows and methods for all data while human-driven analyses would naturally include many variants (using different software, different order in the steps, etc). We argue that the current human-driven processes have resisted creativity and innovation. Each researcher has a tendency to use software that they have used before, since using new algorithms and methods can have a significant learning curve. With NeuroDISK, it would be easy to replace workflow fragments with new methods that become available in the literature. Scientists could easily compare and contrast new methods and see the results of re-running previous analyses using the new methods. We believe our system would stimulate more creativity in terms of exploring new combinations of methods and incorporation of new algorithms.

Intelligent generation of visualizations is also an area of future work. Different types of visualizations may be more useful depending on the scientific question at hand. For example, a visualization may focus on the distribution of the input data for statistical analysis and significance, while others may focus on domain-specific illustrations—like 3D brain images—to aid in exploration. Including these visualizations in our workflows requires domain-specific knowledge from researchers. Visualization is a key aspect for understanding the outcome of an analysis, and easing data exploration may lead to additional scientific questions or prompt researchers to create new datasets to fill in existing gaps.

Because all the scientific questions in NeuroDISK are machine readable, our approach can easily be extended to retrieve semantically similar questions and datasets that have been posed by other investigators and may be related to a new question. NeuroDISK can group similar questions so that their results can be compared. Improving this support for comparing scientific questions may provide researchers with new insights.

NeuroDISK automates many key steps and processes for discovery that are mostly carried out manually today. This is a crucial contribution to the development of AI scientists that can automate scientific exploration and discovery. Today, these steps and processes are typically described as part of the methods section of publications. However, they are not in a machine-readable representation. Moreover, the descriptions of methods are often partial or incomplete(Garijo et al., 2013) which makes it difficult for AI systems to access this information. In contrast, NeuroDISK makes these steps and processes machine-readable and therefore accessible to AI reasoners.

NeuroDISK demonstrates the automation of sophisticated reasoning involved in scientific discovery processes including data retrieval and analysis through planning, reasoning, and execution techniques, leading the way for AI scientists.

Scientific practice would be fundamentally transformed through these developing AI scientists. Scientific advances would be greatly accelerated through the automation of analyses. Pursuing new investigations could potentially be completed in a matter of hours or days rather than months or years. In addition, new results would be more reliable, as they would come with some guarantees of correctness by using lines of inquiry that reflect well known methods and best practices, rather than be prone to human error and misunderstandings. Extensive provenance and explanation would be available for examination, rather than the limited documentation provided in scientific publications. We imagine a future where, once a scientific article is published, its findings are continuously updated when new evidence comes to light, opening the way to fully digital and comprehensively computational paradigms for science.

## 5. Acknowledgments

We are grateful to past and current members of this project for their work on earlier versions of NeuroDISK, particularly Iyad Ba Gari, Joanna Bright, Michael Bornstein, Josh Boyd, Haripriya Dharmala, Vedant Diwanji, Hanna Endrias, Ryan Espiritu, Shobeir Fakhraei, MiHyun Jang, Yibo Ma, Naomi Perez, Abishek Prakash, Wesley Surento, Rosna Thomas, and Regina Wang. We are also grateful to ENIGMA consortium researchers and participants for making this work possible. This work was supported by NIH awards R01AG059874 (PI: Jahanshad) and R01MH134004 (PI: Jahanshad), ONR award N00014-21-1-2437 (PI: Gil), NSF awards IIS-1344272 (PI: Gil) and ICER-1541029 (PI: Gil), and pilot funding from the Kavli Foundation.

## 6. Supplementary Materials / Data Availability

In this section we provide links to web resources, datasets, and workflows developed in our work to date in NeuroDISK and the underlying DISK system:

- Data availability statement: All data and code used in this work are publicly available. The individual cohort results are available as part of the supplementary files of the Grasby et al 2020 publication, and also made available on NeuroDISK. The ABCD dataset, the only one not used in the original Grasby publication, may be accessed at https://nda.nih.gov/abcd. Summary statistics of the genotype-phenotype associations tested here can be accessed through the available NeuroDISK platform.
- DISK:

- DISK software: https://doi.org/10.5281/zenodo.10931904
- DISK system user guide and documentation: https://disk.readthedocs.io/
- DISK’s Scientific Questions Ontology (SQO): https://w3id.org/sqo
- DISK project web site: https://w3id.org/disk
- NeuroDISK:

- NeuroDISK user portal: https://w3id.org/neurodisk
- NeuroDISK Scientific Domain Ontology (SDO): https://w3id.org/enigma
- NeuroDISK data source: https://w3id.org/neurodisk/data
- NeuroDISK project website: https://w3id.org/neurodisk/site

## Notes

https://neuro.disk.isi.edu/

